# Functional dissection of the enhancer repertoire in human embryonic stem cells

**DOI:** 10.1101/146696

**Authors:** Tahsin Stefan Barakat, Florian Halbritter, Man Zhang, André F. Rendeiro, Christoph Bock, Ian Chambers

**Affiliations:** MRC Centre for Regenerative Medicine, Institute for Stem Cell Research, School of Biological Sciences, University of Edinburgh, Edinburgh, EH16 4UU, United Kingdom; CeMM Research Center for Molecular Medicine of the Austrian Academy of Sciences, Lazarettgasse 14, AKH BT 25.3, 1090 Vienna, Austria; Department of Laboratory Medicine, Medical University of Vienna, 1090 Vienna, Austria; Max Planck Institute for Informatics, Saarland Informatics Campus, 66123 Saarbrücken, Germany; Co-first authors; Co-second authors

**Keywords:** ChIP-STARR-seq, genome-wide functional enhancer map, housekeeping enhancers, super-enhancers, transposable elements, naive pluripotency, NANOG, OCT4, H3K27ac, H3K4me1

## Abstract

Enhancers are genetic elements that regulate spatiotemporal gene expression. Enhancer function requires transcription factor (TF) binding and correlates with histone modifications. However, the extent to which TF binding and histone modifications can functionally define active enhancers remains unclear. Here we combine chromatin immunoprecipitation with a massively parallel reporter assay to identify functional enhancers in human embryonic stem cells (hESCs) genome-wide in a quantitative unbiased manner. While active enhancers associate with TFs, only a minority of regions marked by NANOG, OCT4, H3K27ac and H3K4me1 function as enhancers, with activity changing markedly with culture conditions. Our analysis also reveals a novel enhancer set associated with housekeeping genes. Moreover, while transposable elements associate with putative enhancers only some exhibit activity. Similarly, within super-enhancers, large tracts are non-functional, with activity restricted to small sub-domains. This catalogue of validated enhancers provides a valuable resource for further functional dissection of the regulatory genome.

**Highlights:**

- A catalog of functional enhancers in hESCs including a novel housekeeping class
- Active enhancers feature specific transcription factors and transposable elements
- Major shifts in enhancer activity occur during induction of naive pluripotency
- Super-enhancers consist of small units with enhancer function

## Introduction

Human embryonic stem cells (hESC) are a genetically tractable developmental model system with tremendous potential for stem-cell-based therapeutics. Understanding how hESC pluripotency is regulated and how extrinsic signals influence chromatin via transcription factors (TFs) to direct cell-specific gene expression is central to achieving this promise. Gene expression is modulated by *cis*-regulatory elements such as enhancers (Banerji et al., 1981) which can stimulate target gene expression in a position and orientation-independent manner and independent of their genomic context (Spitz and Furlong, 2012). hESCs use a network of pluripotency TFs including OCT4, SOX2 and NANOG, to direct a hESC-specific gene expression programme (Yeo and Ng, 2013). Compared to mouse ESCs, hESCs are a more developmentally advanced state with characteristics of post-implantation stage embryos (Weinberger et al., 2016). Recently, so-called naive human ESCs have been derived from established hESCs either by transient ectopic transgene expression (Buecker et al., 2010; Hanna et al., 2010; Takashima et al., 2014) or by altering culture conditions (Chan et al., 2013; Gafni et al., 2013; Theunissen et al., 2014; Ware et al., 2014). Naive hESCs are thought to mirror cells of the pre-implantation embryo differing from primed hESCs in several ways: increased clonogenicity, different growth factor requirements, distinct energy metabolism, and altered morphology (Sperber et al., 2015). How naïve and primed hESC states are regulated, and how this is affected by differences in enhancer usage is currently not well understood.

The past decade of mammalian genomics research has focused on cataloguing *cis*-regulatory elements within the non-coding genome (ENCODE, 2012). Technological advances have allowed genome-wide occupancy by TFs to be measured by chromatin immunoprecipitation (ChIP) followed by sequencing (ChIP-seq) (Kagey et al., 2010; Robertson et al., 2007). Putative enhancer locations have been obtained by mapping histone modifications (e.g. H3K27ac, H3K4me1) (Heintzman et al., 2007; Rada-Iglesias et al., 2011) and by measuring chromatin accessibility (Buenrostro et al., 2013). However, not all predicted enhancers can be validated functionally. To assay enhancer activity, plasmid-based cell transfections can be used but these have low throughput. More recently, massively parallel reporter assays (MPRAs) have enabled thousands of sequences to be tested simultaneously (Arnold et al., 2013; Kwasnieski et al., 2012; Melnikov et al., 2012; Patwardhan et al., 2012; Smith et al., 2013). For instance, with Self-Transcribing Active Regulatory Region Sequencing (STARR-seq) compact, non-mammalian genomes can be screened quantitatively for enhancer activity by cloning randomly sheared DNA between a minimal-promoter-driven GFP open reading frame and a downstream polyA sequence. If an enhancer is active, this results in transcription of the enhancer sequence (Arnold et al., 2014; Arnold et al., 2013; Shlyueva et al., 2014). Similar MPRA approaches have recently been adapted to test chosen sequences with putative enhancer features (Kwasnieski et al., 2014; Shen et al., 2016; Vanhille et al., 2015), predicted TF binding sites (Verfaillie et al., 2016), features of quantitative trait loci (Tewhey et al., 2016; Ulirsch et al., 2016; Vockley et al., 2015) or nucleosome-depleted sequences (Murtha et al., 2014).

Application of STARR-seq to explore mammalian genomes is hindered by genome size which means enhancer sequences would be infrequently sampled and transfection of plasmid libraries would require huge numbers of cells. Here we alleviate this issue by combining STARR-seq with ChIP, in a technique we refer to as ChIP-STARR-seq to generate a resource of genome-wide activity maps of functional enhancers in hESCs. In these maps we identify highly active enhancers and observe major changes in activity patterns between primed and naive hESCs. Moreover, some transposable element families are enriched at highly active enhancers. Our data also identify the functional components within super-enhancers and uncover a novel class of enhancers associated with housekeeping gene expression. The resource presented here encompasses a comprehensive collection of functional enhancer sequences in hESCs, providing a valuable knowledge base for systematic analysis of the core transcriptional network circuitry underlying hESC maintenance and differentiation. Enhancer data are available from the Supplemental Materials to this paper and from a supplemental website (http://hesc-enhancers.computational-epigenetics.org).

## Results

### ChIP-STARR-seq: an effective strategy for genome-wide identification of functional enhancers

To generate a comprehensive catalogue of regulatory genomic elements relevant to mammalian stem cell biology we used a massively parallel reporter assay, called ChIP-STARR-seq. In ChIP-STARR-seq, DNA is co-immunoprecipitated and cloned *en masse* within the transcription unit of a STARR-seq plasmid, downstream of GFP driven by a minimal promoter and upstream of a polyA sequence (Figure 1A)(Arnold et al., 2013). The resulting libraries, each consisting of millions of individual plasmids containing DNA sequences that were bound by a factor of interest, can be tested for enhancer activity by cell transfection. If a cloned sequence functions as an enhancer, GFP will be expressed and the transfected cells can be purified by FACS. Since the assayed sequences are located upstream of the polyA signal, the transcribed mRNA will contain the enhancer sequence itself. Therefore, both the identity and activity of captured regions can be determined quantitatively by sequencing mRNA (RNA-seq) from GFP-positive cells.

**Figure 1:**
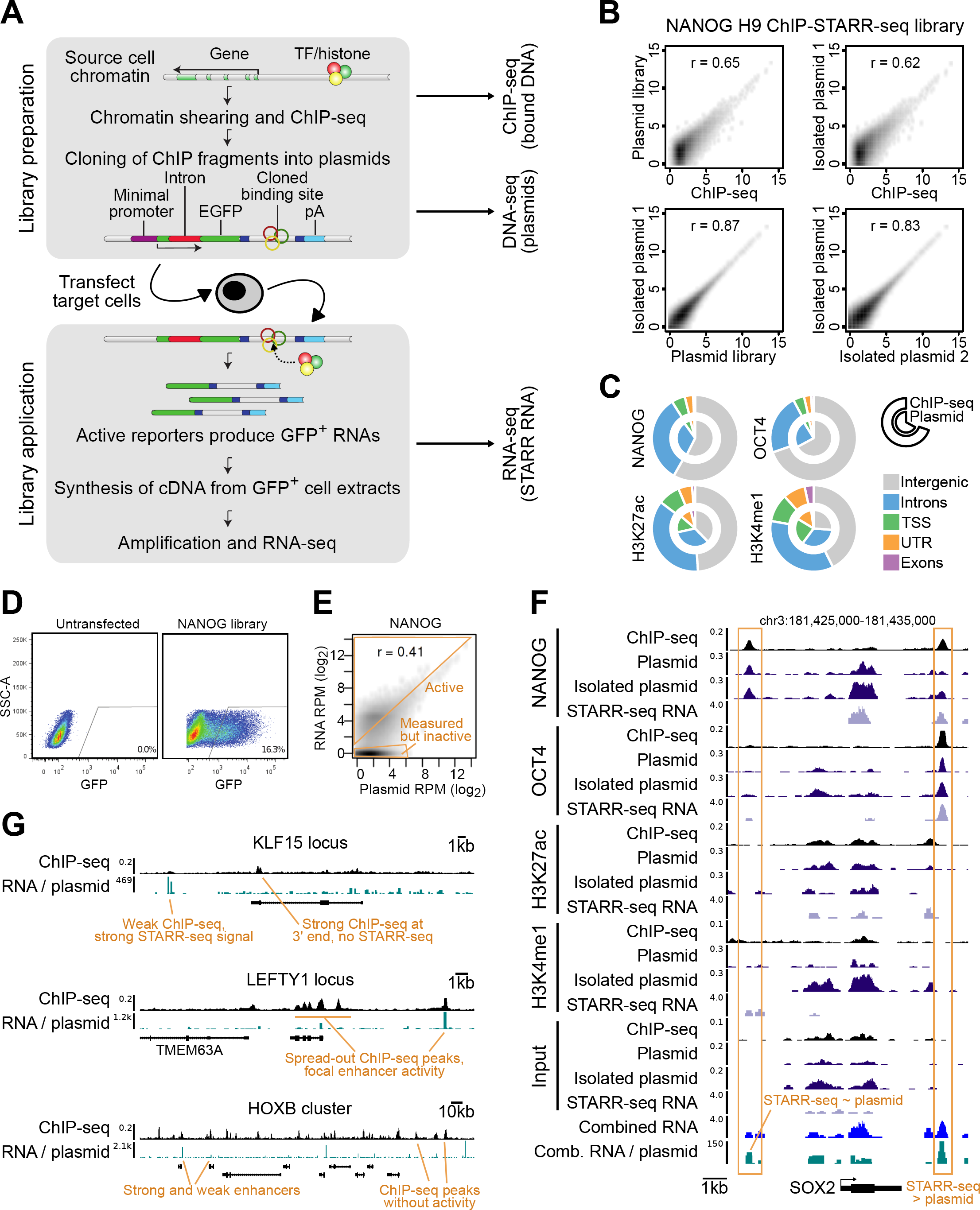
ChIP-STARR-seq in human embryonic stem cells. **A**) To prepare ChIP-STARR-seq plasmid libraries, ESC chromatin is sonicated to between 200-600bp and immunoprecipitated with antibodies against TFs or histone modifications (coloured balls). DNA is end-repaired, adapter-ligated and used to make a plasmid library by Gibson assembly. In the STARR-seq plasmid (Arnold et al., 2013) a multiple cloning site (MCS; dark blue) lies downstream from a minimal promoter (purple) and an intron-containing EGFP ORF (green) and upstream from a UTR/polyA sequence (light blue). Plasmids libraries are then transfected into target cells. If cloned DNA has enhancer activity, transcription from the minimal promoter is activated, yielding GFP mRNAs with enhancer sequences embedded in the 3’UTR. Enhancers can then be identified by FACS purification of GFP-positive cells and RNAseq. **B**)ChIP-STARR-seq for NANOG in H9 hESCs. Each scatterplot contrasts the normalized read count (reads per million) per peak between two datasets, obtained from ChIP-seq or DNA-seq of the generated plasmid libraries before or after transfection and recovery from hESCs (n=2). The Pearson correlation (*r*) for each comparison is indicated. **C**) Doughnut charts of the genomic distribution of peaks called for ChIP-seq (outer chart) and corresponding plasmid libraries (inner chart); TSS, transcription start sites; UTR, untranslated region. **D**) FACS plots of single, DAPI-negative hESCs. Left, untransfected cells; right, cells transfected with a NANOG ChIP-STARR-seq plasmid library. **E**) Scatterplot (as in panel B) contrasting the NANOG plasmid library and corresponding ChIP-STARR-seq RNA. The dense cluster of points in the lower left corner corresponds to library plasmids that did not produce RNAs. RPM, reads per million. **F**)Genome browser plot of the *SOX2* locus showing tracks for ChIP-seq, DNA-seq of plasmid libraries, and ChIP-STARR-seq from RNA-seq of GFP^+^ cells transfected with the indicated libraries. **G**) Genome browser shots of the *KLF15* and *LEFTY* loci and *HOXB* cluster, illustrating a broad variety of enhancers profiled in this catalogue of functional enhancers.

To investigate the functional potential of enhancers in hESCs, we first focused on H9 hESCs cultured on Matrigel, in medium supplemented with b-FGF. In these primed culture conditions hESCs grew as flat colonies expressing NANOG and OCT4 (Figure S1A, B). ChIP-qPCR confirmed enrichment of sequences known to be bound by NANOG, OCT4, H3K4me1 and H3K27ac in hESCs (Figure S1C) (Kunarso et al., 2010; Rada-Iglesias et al., 2011). Subsequent ChIP-seq confirmed a high overlap with peaks from previous datasets (Figure S1D) (Gafni et al., 2013; Gifford et al., 2013; Ji et al., 2016; Kunarso et al., 2010; Lister et al., 2009).

We generated ChIP-STARR-seq libraries from the same end-repaired, adapter-ligated ChIP DNA used for the ChIP-seq experiments (Figure 1A). Total genomic DNA was also used to carry out STARR-seq in hESCs (Arnold et al., 2013). To generate reporter plasmid libraries, we cloned ChIP DNA *en masse* into the STARR-seq plasmid by Gibson assembly. Sequencing the resulting plasmid libraries produced 1.9x10^9^ reads. Counting unique read ends mapping to the human genome, indicated that each library consisted of 1.9-3.1x10^7^ unique plasmids, with a mean insert size of 230bp (**Table S1**). (Figure S2A) summarises the sequenced samples analysed in this study.

To validate the comprehensiveness of our reporter assay, we first assessed whether the plasmid libraries achieved a good representation of the binding events captured by ChIP-seq (**File S1**). A high correlation between ChIP-seq coverage and the corresponding plasmid libraries was seen both before and after transfection (Figure 1B,C S2B,C). Next, the ability of the plasmid libraries to drive GFP expression in primed hESCs was tested. Library transfections produced up to 20% GFP-positive cells compared to <1% GFP-positive cells obtained by transfection of the empty STARR-seq vector or ∼50% in control transfections with a constitutively expressed mCherry plasmid (Figure 1D and data not shown). Therefore, a considerable proportion of cells contained plasmids with enhancer activity. 24h after transfection RNA was prepared from FACS-purified GFP-positive cells and DNA from unsorted cells. Plasmids were recovered and GFP-derived mRNAs amplified by PCR for RNA-seq. DNA sequencing confirmed high consistency between the original plasmid libraries and plasmids re-isolated after transfection (Figure 1B, S2C). Positive correlations were also observed between read coverage from STARR-RNA-seq and the respective plasmid libraries (Figure 1E, S2D) and between replicate STARR-RNA-seq datasets, with an increase for expressed plasmids sampled (read count > 0) in both replicates (Figure S2E). Taken together, these results show that while abundant plasmids can produce more RNA, some plasmids produce RNA in excess of the plasmid count, indicating high enhancer activity. However, many plasmids transfected into cells did not produce RNA indicating that the ChIP-enriched DNA in these plasmids lacked enhancer activity.

Visual inspection of selected genomic regions illustrates the broad spectrum of enhancer activity measured by ChIP-STARR-seq (Figure 1F,G). For instance, ChIP-seq for NANOG indicates two strong binding sites upstream and downstream of *SOX2* (Figure 1F) but only the downstream binding site resulted in ChIP-STARR-seq RNA in excess of plasmid abundance.

In summary, we used ChIP-STARR-seq for a functional enhancer analysis of DNA fragments bound by NANOG, OCT4, H3K4me1, and H3K27ac in hESCs.

### Activity levels define classes of enhancers bound by distinct transcription factors

Based on our ChIP-STARR-seq dataset, we assessed the functional capacity of 296,034 genomic regions represented in our plasmid libraries. Enhancer activity was defined as the ratio of RNA reads relative to plasmid reads scaled to account for differences in sequencing library size (RPPM: reads per plasmid and per million sequenced reads). Paired-end sequencing enabled unequivocal assignment of RNA reads to plasmids. The activity level of each region was recorded as the activity generated by the most active plasmid (from any library) within this region. Thresholds for discriminating enhancer activity from the activity of the minimal promoter in the STARR-seq vector were obtained by comparison to data from inactive regions (see Methods). This calculation defined a threshold for high and low activity elements of ≥220 and ≥144 RPPM, respectively. Based on these thresholds, ChIP-STARR-seq identified 21,200 high activity enhancers and 18,422 low activity enhancers (Figure 2A, **File S1**). However, the majority of peaks showed no evidence of enhancer activity (Figure 2A,B).

**Figure 2:**
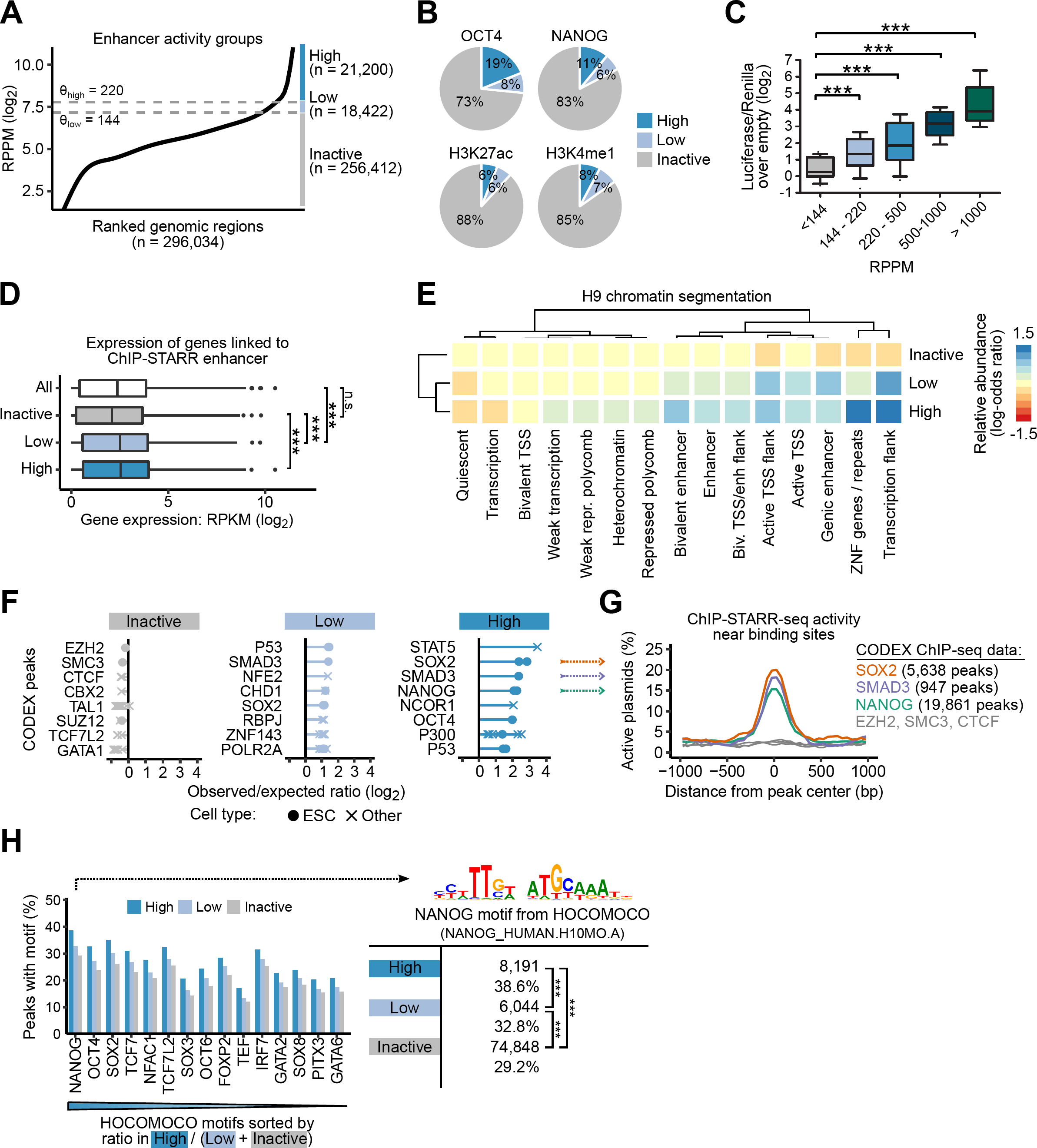
Activity levels define functional classes of enhancers. **A**)Plot showing enhancer activity (enrichment of ChIP-STARR-seq RNA over plasmids; log_2_) ranked from lowest to highest across all measured enhancers (union of all peak calls). Three groups of enhancers were distinguished based on their activity and the thresholds (θ) are indicated by dashed lines. RPPM, reads per plasmid and per million sequenced reads. **B**)Distribution of high activity (RPPM ≥220), low activity (RPPM 144-220) and inactive sequences (RPPM <144) in peaks called for the indicated factors. **C**)Luciferase activities of 68 tested genomic sequences in primed hESCs grouped by ChIP-STARR-seq activity. Activity is the fold enrichment relative to empty vector (in log_2_), normalised to a Renilla transfection control. Boxes are interquartile range, line is median, whiskers are 10^th^ to 90^th^ percentile. ***= p<0.001, unpaired t-test; n=2. **D**)Distribution of gene expression values (Takashima et al., 2014) of genes associated with enhancers grouped by activity level. Boxplots represent interquartile range (IQR), line is the median, whiskers extend to 1.5xIQR, dots indicate outliers. All, all genes in RNA-seq dataset; inactive, RPPM <144; low, RPPM 144-220; high, RPPM ≥220; RPKM, reads per kilobase million. **E**) Heatmap of the relative enrichment of H9 chromatin segments from the Roadmap project (Kundaje et al., 2015) in regions with different ChIP-STARR-seq activities (see panel A). Colors report log-odds ratios. Rows and columns are arranged by hierarchical clustering with complete linkage to put similar segment types together. TSS, transcription start site; enh, enhancer; ZNF, zinc-finger protein. **F**)Relative enrichment of TFs from the CODEX database (Sanchez-Castillo et al., 2015) in inactive regions, low activity and high activity enhancers. Shown are the log_2_-odds ratios between observed percentages of enhancers overlapping binding sites for each TF in the respective groups relative to the percentage in the entire region set. Each dot represents a TF ChIP-seq dataset with lines connecting the most extreme dot to zero for visualization. For each category, the eight most enriched TFs are ranked by their mean log-odds ratio. ChIP-seq datasets from hESCs are indicated as dots and those from other cells as crosses. Enrichments were calculated using LOLA (Sheffield and Bock, 2016). **G**)Smooth line plots showing proportion of active plasmids (RPPM ≥220) of all plasmids measured at the indicated distance from the peak center for ChIP-seq binding sites for SOX2, SMAD3 and NANOG averaged across all binding sites for the respective factor. **H**) Top motifs from the HOCOMOCO database (Kulakovskiy et al., 2016) ranked by the hit frequency within the high activity enhancers compared to low and inactive enhancers (strongest enrichment is at the left). Numbers indicate the proportion of peaks with ≥1 motif hit. Motif matches were determined using FIMO (Grant et al., 2011). On the right, the DNA sequence logo of the top motif is shown. ***, p < 0.001, Fisher’s exact test.

To assess the biological relevance of these thresholds, sixty-eight genomic regions covering the full activity range were tested in luciferase assays. DNAs from regions of <144 RPPM had luciferase activities indistinguishable from empty vector. In contrast, regions with increasingly high ChIP-STARR-seq activity showed increasingly higher luciferase activity (Figure 2C) supporting the utility of the RPPM-based enhancer classification. These thresholds indicate that only a minority of peaks bound by NANOG, OCT4, H3K4me1 or H3K27ac showed enhancer activity (Figure 2B), with regions bound by OCT4 having the highest proportion of high activity enhancers.

To assess the relationship of activity classifications to gene expression, each peak was assigned to a putative target gene based on genomic distance (Figure 2D, S3B). ChIP-STARR-seq regions with enhancer activity were associated with genes that showed significantly higher gene expression values than genes associated with peaks lacking enhancer activity. Regions with different ChIP-STARR-seq activity levels were next assessed for association with distinct histone modifications in an H9 chromatin segmentation (Kundaje et al., 2015) (Figure 2E). Chromatin segments marked as enhancers, transcription start sites (TSSs), sites flanking transcription and repeat sequences were most overrepresented in the high activity group. Together, these results indicate that ChIP-STARR-seq can distinguish ChIP-seq peaks on the basis of enhancer activity and this enhancer activity reflects expression of the endogenous sequences.

The occurrence of TF binding sites in the three activity classes was next assessed. The relative representation of TFs from 190 ChIP-seq datasets in the CODEX database were assessed by LOLA enrichment analysis (Sanchez-Castillo et al., 2015; Sheffield and Bock, 2016) (Figure 2F, Table S2). High activity enhancers were preferentially associated with pluripotency-related TFs (SOX2, SMAD3 and NANOG). Significant overlaps were also seen for regions bound in non-hESCs by STAT5 and NCOR1. Low activity enhancers were weakly enriched for pluripotency-related TFs (SOX2 and SMAD3) but also for proteins with more generic functions (NFE2, CHD1, RBPJ, ZNF143). In contrast, no TF in the reference database were enriched at inactive regions. Similar results were obtained by extending LOLA analysis to 690 ChIP-seq datasets for TFs from ENCODE (2012) (Figure S3D). Enhancer activity was strongest close to the binding peaks of the enriched factors with activity lost quickly with increasing distance from the peak center (Figure 2G, S3C). These results suggest that binding of distinct TFs in close proximity might contribute to robust enhancer activity. How enhancer classes relate to chromatin state was further examined by LOLA analysis of ENCODE chromatin segmentations from H1 hESCs and various non-pluripotent cell types (Figures S3E,F). This confirmed that high activity enhancers were enriched in segments annotated as H1 enhancers and promoters, while inactive regions occurred primarily in closed chromatin. Interestingly, low activity enhancers were often annotated in ENCODE as non-cell-specific promoters. We also interrogated the DNA sequences underlying enhancers for the occurrence of known TF binding motifs (Figures 2H, S3G). The most enriched motif in the highly active enhancers was the designated NANOG motif in the reference database, which we note actually corresponds closely to the Oct/Sox motif from the literature (Chen et al., 2008). The percentage of peaks that contained this motif was higher in the high activity group. These results indicate that functional classes of enhancers differ in TF binding sites and motif occurrence.

### Active enhancers include ESC-specific and housekeeping modules

Previous high-throughput sequencing studies have attempted to predict hESC enhancers on the basis of histone marks, TF binding sites or DNAseI hypersensitivity (Hawkins et al., 2011; Rada-Iglesias et al., 2011; Xie et al., 2013). However, the overlap between enhancers predicted from these studies is limited (Figure S4A). We sought to exploit the catalogue of functional enhancers to ratify enhancer predictions. Comparing the combination of three previously described enhancer maps with our dataset, 7,596 of the 21,200 high activity enhancers identified by ChIP-STARR-seq were among these predicted enhancers (n = 76,666; union of all datasets; **Table S3**). Several putative enhancers predicted by these previous studies that were inactive by ChIP-STARR-seq were tested in luciferase assays but none possessed enhancer activity in this assay (Figure S4C). Functional enrichment analysis using GREAT (McLean et al., 2010) showed that the high activity ChIP-STARR-seq enhancer subset overlapping with previously predicted enhancers had stronger enrichment for gene ontology (GO) terms related to ESC biology than terms identified from all predicted enhancers (**Table S2**). This “ESC module” (Figure 3A) includes enhancers in close proximity to hESC TFs (*NANOG, OCT4)* and signaling pathway genes (TGF-b, FGF, and WNT signaling). The remaining 13,604 enhancers with high ChIP-STARR-seq activity, that were not predicted previously, had GO terms reflecting generic biological processes, for instance, transcription, energy metabolism and DNA repair. Hence, we refer to these enhancers as the “housekeeping (HK) module”.

**Figure 3:**
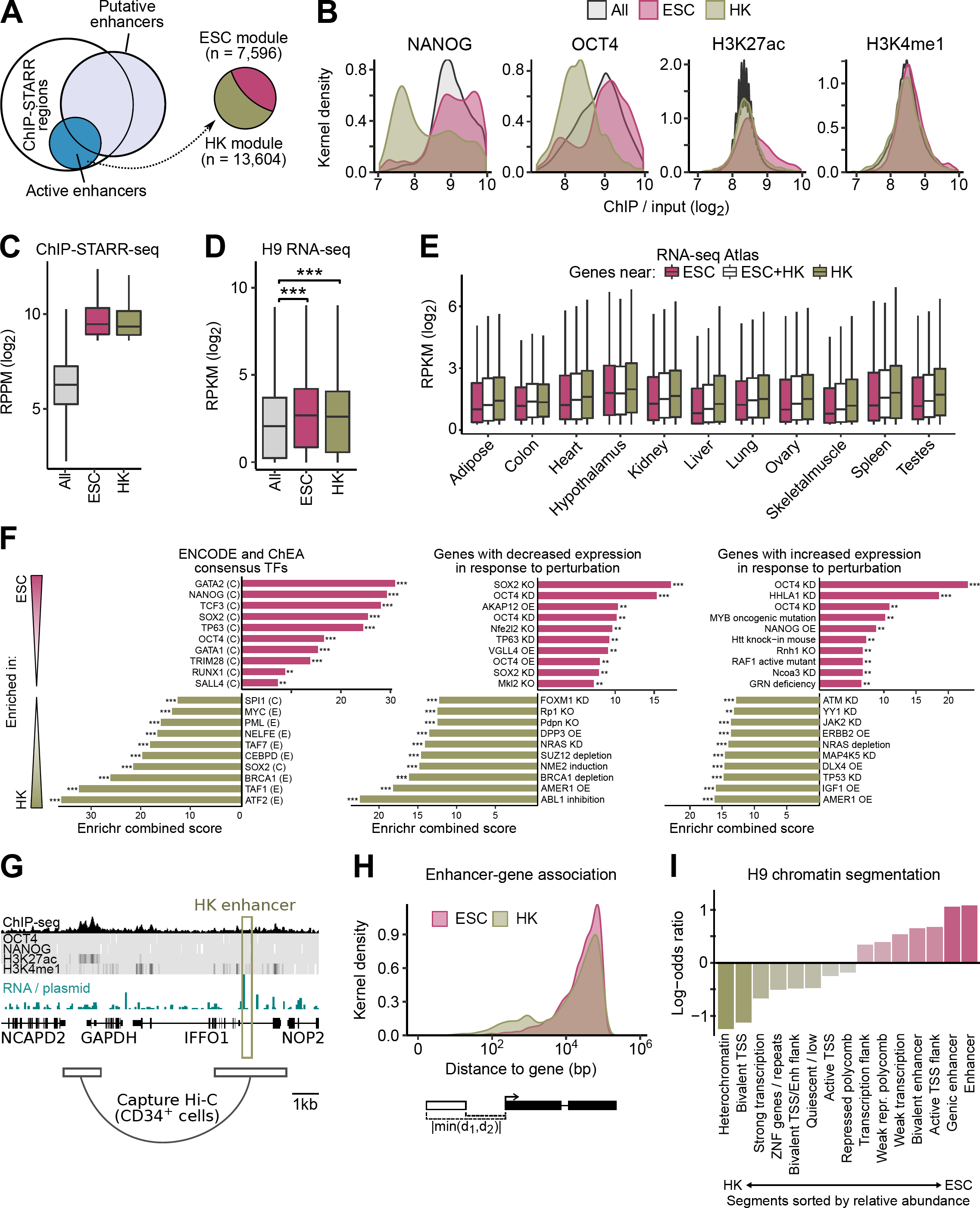
Active enhancers include ESC-specific and housekeeping modules. **A**)Illustration of the overlap between previously published putative enhancers (Hawkins et al., 2011; Rada-Iglesias et al., 2011; Xie et al., 2013) (light blue circle) and regions assessed in ChIP-STARR-seq in hESCs (white circle). Highly active ChIP-STARR-seq regions (RPPM ≥220) are indicated by the blue circle. ChIP-STARR-seq enhancers overlapping with previously published putative enhancers are referred to as the ESC module and non-overlapping ChIP-STARR-seq enhancers as the housekeeping (HK) module. **B**) Kernel density plots of the distribution of enrichment values (ratio of ChIP-seq signal over input, log_2_) in hESCs for NANOG, OCT4, H3K27ac and H3K4me1 for all peaks called or for peaks associated with ESC or HK module enhancers. **C**) Distribution of measured RPPM values for all assessed genomic regions compared to enhancers from the ESC and HK modules. Boxplots represent interquartile range (IQR), line is the median, whiskers extend to 1.5xIQR. RPPM, reads per plasmid million. **D** Gene expression (RNA-Seq; log_2_) in H9 hESCs (Takashima et al., 2014) for all genes compared to genes associated with either ESC or HK module enhancers. Boxplots as in C. RPKM, reads per kilobase million. *** = p<0.001 (t-test). **E**) Gene expression (RNA-Seq; log_2_) in different tissues from the RNA-seq Atlas (Krupp et al., 2012) for all genes linked to ESC or HK module enhancers or both. Boxplots as in C. **F**) Functional enrichment analysis using Enrichr testing the relative over-representation in ESC (top) and HK (bottom) module enhancers near genes associated with different annotation categories (Chen et al., 2013). Shown are the top 10 results each for TF binding sites from ENCODE and ChEA (left) and genes down-regulated (middle) or up-regulated (right) upon single-gene perturbations from the GEO database. The x-axis reports the combined score calculated by Enrichr. **G**) Genome browser of *GAPDH* locus including neighbouring genes. The olive-green box highlights one HK enhancer with high activity. Chromosome confirmation data from other cell types (Mifsud et al., 2015) indicates that this region interacts with the promoter of *GAPDH*. **H**) Kernel density plot showing the distance to associated genes for ESC and HK module enhancers. Distance was calculated between both boundaries of the enhancer region and the start site of the linked gene with the shorter absolute distance recorded. **I**) Bar plot indicating the relative enrichment (log-odds ratio in ESCs compared to all) of H9 chromatin segments from the Roadmap Epigenomics project (Kundaje et al., 2015) in ESC and HK enhancers. Segment types more commonly overlapping with HK modules (left) or ESC enhancers (right) are shown.

A comparison of the ChIP-seq signal intensity for all peaks, or peaks associated with either the ESC or HK modules, indicates that HK enhancers generally, had lower affinity for H3K4me1, H3K27ac, NANOG and OCT4 (Figure 3B). Therefore, these HK enhancers may have escaped detection previously due to the thresholds used. Applying instead a direct functional assay enabled their discovery. The lower levels of NANOG and OCT4 at the HK enhancers may suggest that these enhancer sequences rely less on ESC-specific TFs. Nonetheless, these sequences function as *bona fide* enhancers, as the RPPM of these sequences is higher than those of all sequences assessed (Figure 3C) and only slightly lower than those seen in the ESC module. In addition, the expression of genes associated with the ESC and HK module is similar and significantly above the average hESC gene expression levels (Figure 3D).

Consistent with function in many cell types, expression of genes associated with the HK module was higher than expression of genes associated with the ESC module in data from various tissues of the RNA-seq Atlas (Krupp et al., 2012) (Figure 3E)and from the GTEx portal (2013) (Figure S4B,D). Furthermore, functional enrichment analysis using the Enrichr software (Chen et al., 2013) with data from ENCODE (2012) or ChEA (Lachmann et al., 2010) showed that ESC module enhancers were enriched for binding of NANOG, TCF3, SOX2 and OCT4, whereas HK module enhancers showed preferential enrichment of more broadly expressed factors, such as BRCA1 and MYC (Figure 3F, Table S2). For instance, one of the HK enhancers that we identified is located upstream of *GAPDH*, in a region that has been shown to interact with the *GAPDH* promoter in various cell types (Figure 3G). The majority of HK and ESC enhancers showed a similar distribution of distances from TSSs, although a subset of HK enhancers lie closer to TSSs (Figure 3H). ESC enhancers were often found in regions associated with enhancer-like chromatin features in H9 hESCs (Kundaje et al., 2015) (Figure 3I). In contrast, HK enhancers were more often annotated as heterochromatic or bivalent. ChIP-STARR-seq therefore identified previously unappreciated genomic sequences characterized by lower enrichment of enhancer-associated histone modifications and pluripotency-related TFs but with comparable enhancer activity.

### Major changes in enhancer activity upon induction of naive pluripotency

To augment the catalogue of functional enhancers in hESCs and to gauge the dynamics of enhancer activity we applied ChIP-STARR-seq to a closely related cell type. To this end, primed H9 hESCs were passaged in Naive Human Stem Cell (NHSM) medium to establish phenotypically altered, “naïve” hESCs (Gafni et al., 2013). Dome-shaped colonies expressing NANOG and OCT4 appeared after 3 days that could be passaged more than 10 times (Figure S5A, B). Relative to primed hESCs these naive hESCs expressed similar levels of *OCT4*, *REX1* and *STAT3* but with higher *NANOG*, *TEAD4*, *KLF4*, *DUSP10*, *IL6R, TBX3* and lower *XIST* and *DNMT3B* (Figure S5C,D). ChIP-qPCR showed that selected loci were similarly bound by NANOG, OCT4, H3K4me1 and H3K27ac in both primed and naive hESCs (Figure S5E). A high overlap of H3K4me1 and H3K27ac ChIP-seq peaks with previous data was seen (Gafni et al., 2013) (Figure S5F,G). NANOG and OCT4 ChIP-seq data identified similar motifs in both naive and primed hESCs (Figure S5H,I). These results agree with prior studies (Barakat et al., 2015; Gafni et al., 2013) confirming conversion of primed hESCs to a naive hESC state.

ChIP-STARR-seq plasmid libraries generated from naive hESCs (Figures 4A, S6A-C) were transfected into naive hESCs and for comparison, into primed hESCs (Figure S6C). Enhancer activity was categorized with the same calculation as in primed hESCs into high activity (RPPM ≥96) and low activity (RPPM ≥62) (Figure S6D, S2E, **File S1**). 337,178 peaks covered by plasmids in naive hESCs (Figure S6E) were analysed, identifying 31,303 high activity and 40,732 low activity enhancers. Again, only a fraction of ChIP-seq peaks had high activity (Figure S6F). LOLA enrichment analysis of TFs from CODEX for the naive enhancer class (Figure 4B, Table S2), identified a distinct TF profile from primed hESCs (compare to Figure 2F). Sites bound by pluripotency-related TFs (e.g., NANOG) in primed hESCs were not strongly represented in the high activity enhancer category. Instead, genomic regions bound by repressive chromatin interactors (CHD1, HDAC1/2) in primed hESCs showed high activity in naive hESCs. The altered protein binding landscape was also seen by enrichment analysis of ENCODE ChIP-seq datasets (Figure S6G) and chromatin segmentations (Figure S6H) (Ernst et al., 2011). Enhancers with high activity in naive cells occurred mainly in regions that, in primed cells, were bound by TFs linked to proliferation and often to cancer (BRCA1, FOSL1, MYC). These enhancers overlap with active promoters in various cell types more than with primed enhancers (Figure S6H, I).

**Figure 4:**
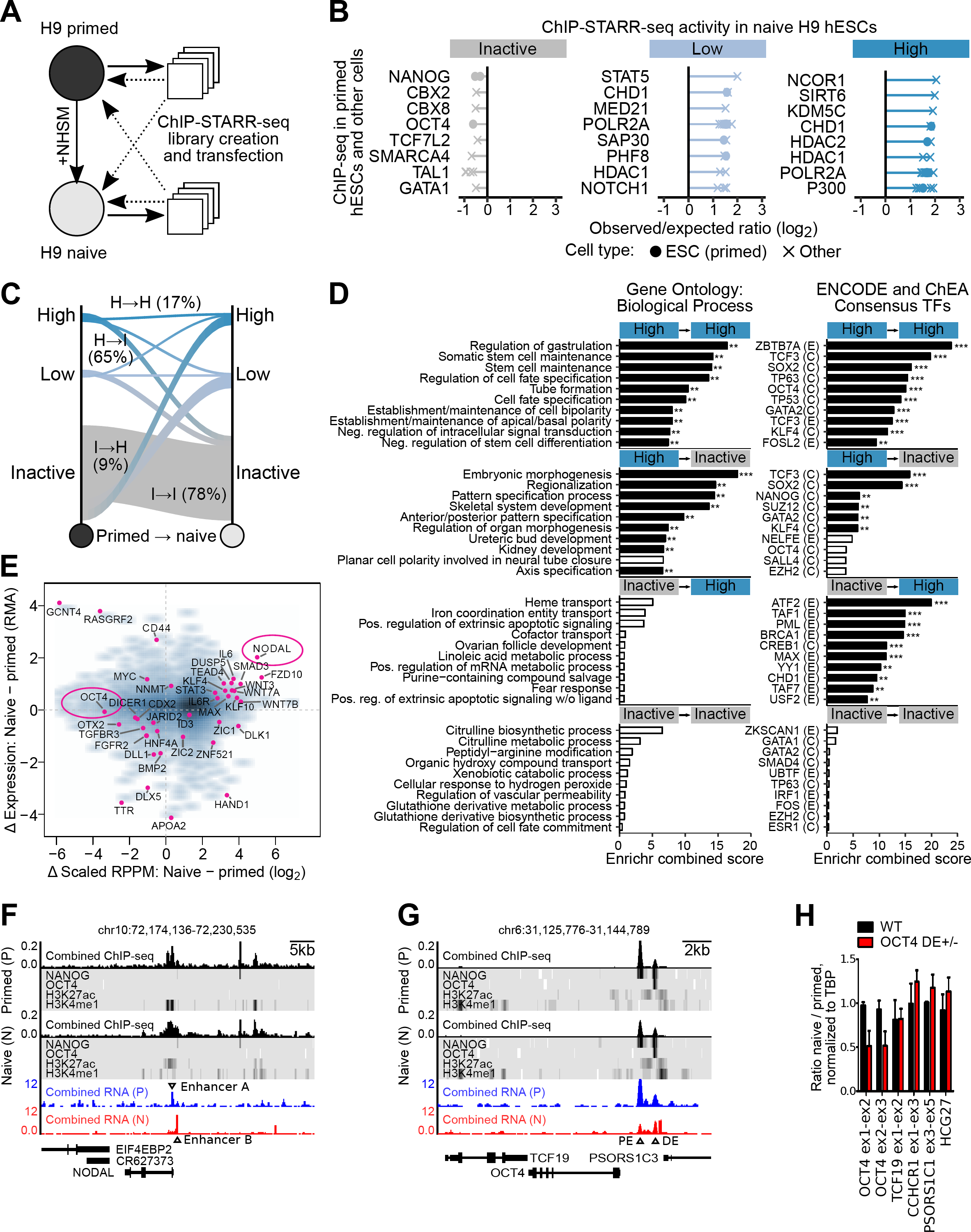
Major changes in enhancer activity upon induction of naive pluripotency. **A**)Overview of the primed to naive conversion and ChIP-STARR-seq cross-over design. **B**)Relative enrichment of TFs from CODEX (Sanchez-Castillo et al., 2015) in inactive, low activity and high activity enhancers in naive H9 hESCs. Plots as in Figure 2F. ChIP-seq datasets produced from primed hESCs are indicated as dots and those from other cell sources as crosses. **C**) Sankey river plot illustrating relative changes in enhancer activity between primed (left) and naive (right) hESCs. Percentages refer to the total number of peaks of the respective group in primed hESCs. Percentages and groups are indicated only for groups with the most prominent changes or those that maintain activity. **D**) Functional enrichment analysis using Enrichr to test the relative over-representation of enhancers near genes with certain GO assignments (left) or occurring near binding sites from ENCODE and ChEA ChIP-seq experiments (right). Shown are the top 10 results for each enhancer group highlighted in panel F. The x-axis reports the combined score calculated by Enrichr. **E**) Scatterplot contrasting changes in enhancer activity with changes in gene expression of associated genes. RPPM values are rescaled by dividing them by the cell-state-specific high-activity threshold prior to calculating the difference between primed and naive hESCs. Outliers and assorted genes are highlighted with dots and examples are labeled. **F**)Genome browser shot of *NODAL*. ChIP-seq tracks for NANOG, OCT4. H3K27ac, and H3K4me1 in primed (top) and naive (middle) hESCs are shown, together with the ChIP-STARR-seq RNA (combination of all datasets) in the respective cell states. A switch of enhancer activity from the promoter-overlapping enhancer A to the upstream enhancer B is observed in the primed to naive transition. **G**) Genome browser shot of the *OCT4* locus. Display equivalent to E. The proximal enhancer (PE) and distal enhancer (DE) display different activity patterns in naive and primed hESCs. **H**) qRT-PCR analysis of wild type (wt) H9 hESCs or H9 hESCs with a heterozygous *OCT4* distal enhancer deletion (DE+/-) for two amplicons detecting OCT4 mRNA or mRNAs from flanking genes. The average ratio of gene expression in cells cultured in naive conditions (8 passages) relative to primed conditions is shown for three wt or three *OCT4* ΔDE+/- cell lines, normalized for TBP and one wt set to 1. * =p<0.05 (2-way ANOVA with Bonferoni post-test), error bars are SD.

Having a comprehensive genome-wide enhancer maps for both pluripotent states allowed a global comparison of enhancer usage in both primed and naive hESCs (Figure 4C). We focused on regions that maintained (H→H) or lost high enhancer activity (H→I), that remained inactive (I→I) or that gained high activity (I→H) in the primed to naive switch. Functional enrichment analysis of these four groups with Enrichr (Figure 4D, Table S2) revealed that enhancers with high activity in both cell states (H→H) were related to stem cell maintenance and suppression of differentiation, whereas enhancers that lost activity (H→I) were associated with genes with GO terms related to differentiation. No significant GO terms were associated with enhancers that gained activity (I→H) or regions that remained inactive (I→I), though this may be due to lack of annotation in naive hESCs. However, examining ENCODE and ChEA indicated that enhancers that gained activity in naive hESCs were enriched for transcriptional activators such as ATF2 and TAF1 that occur near target gene promoters.

To relate changes in enhancer activity to differences in the expression of regulated genes, we plotted the average difference in enhancer RPPM levels between naive and primed hESCs versus the expression of nearby genes (Figure 4E). Several genes previously found to be more strongly expressed in naive hESCs (such as members of the WNT pathway) showed increased enhancer activity and vice versa.

To further compare enhancer activity in primed and naive hESCs, two loci were examined in more detail (Figure 4F). Two adjacent enhancers proximal to the *NODAL* promoter exhibit a pattern typical of the changing activity landscape (Figure 4F). In primed hESCs, enhancer A is highly active, yet the *NODAL* gene is weakly expressed. Enhancer A is marked by H3K4me1 but lacks H3K27ac suggesting that site A may function as an enhancer rather than as the promoter of the adjacent gene. In naive hESCs, enhancer A activity is lost but enhancer B is activated. Concordantly, NODAL expression is elevated and an increase in H3K27ac intensity is observed at the promoter.

At *OCT4,* a similar binding of NANOG to the proximal (PE) and distal (DE) enhancers was seen in primed and naïve hESCs (Figure 4G). However, in primed hESCs, ChIP-STARR-seq activity was mainly detected from the PE. In naïve hESCs, PE activity was strongly reduced, whereas DE activity was similar in both cell states (Figure 4G). Independent luciferase assays confirmed these findings (Figure S7D). To determine the biological relevance of this switch in enhancer usage, primed hESCs with heterozygous OCT4 DE deletions were generated by CRISPR-Cas9 (Figure S7). These primed OCT4 ΔDE+/- hESCs expressed similar levels of *OCT4* mRNA to wild type clones. However, *OCT4* mRNA dropped to 50% upon conversion to the naive state without affecting flanking gene expression (Figure 4H). This indicates that OCT4 DE is indeed a functional enhancer that regulates OCT4 in naive hESCs. We conclude that enhancer activity is remarkably dynamic even in closely related cell types.

### The occurrence of various transposable elements is associated with enhancer activity

As chromatin segments associated with repetitive DNA were found in high activity enhancers (Figure 2E), we sought to exploit the functional enhancer catalogue presented here to examine the link between repeats and enhancer activity more closely. Large portions of mammalian genomes are derived from transposable elements (TEs) which have been reported to be linked to TF binding sites and enhancers (Bourque et al., 2008; Glinsky, 2015; Kunarso et al., 2010; Teng et al., 2011). In hESC, human endogenous retrovirus (HERV) TEs are enriched in NANOG and OCT4 binding sites (Glinsky, 2015; Kunarso et al., 2010) but whether this enrichment reflects enhancer activity has not been determined genome-wide. To assess ChIP-STARR-seq enhancers for the occurrence of TE sequences, we used the RepeatMasker annotation in the UCSC Genome Browser. The number of TE-derived sequences in regions of distinct activity was compared to the number detected in all genomic regions (Figure 5, Table S4). LTR-containing TEs, such as HERV1, were found in high activity enhancers more often than expected (Figure 5A). However, not all LTR-containing TEs were enriched at active enhancers. The most enriched repeats were dominated by satellite repeats and LTR family members (Figure 5B). For TEs enriched for NANOG and OCT4 binding (e.g., LTR9B) (Kunarso et al., 2010) or TEs enriched at candidate human-specific regulatory loci (e.g., LTR7) (Glinsky, 2015) the observed enrichment increases further with increasing enhancer activity (Figure 5C, D). Indeed LTR7B, LTR7 and HERVH-int show the strongest enrichment at the highest activity enhancers. In contrast, other TE families that have been previously linked to human-specific TF binding sites (Glinsky, 2015), were either not (L1HS) or only weakly (L1PA2) enriched at high activity enhancers. These results indicate that certain families of TEs are overrepresented at active enhancers and that their enrichment correlates with enhancer activity However, not all TEs of the same TE type are associated with active enhancers, nor do all TEs enriched in pluripotency TF binding sites occupy active enhancers.

**Figure 5:**
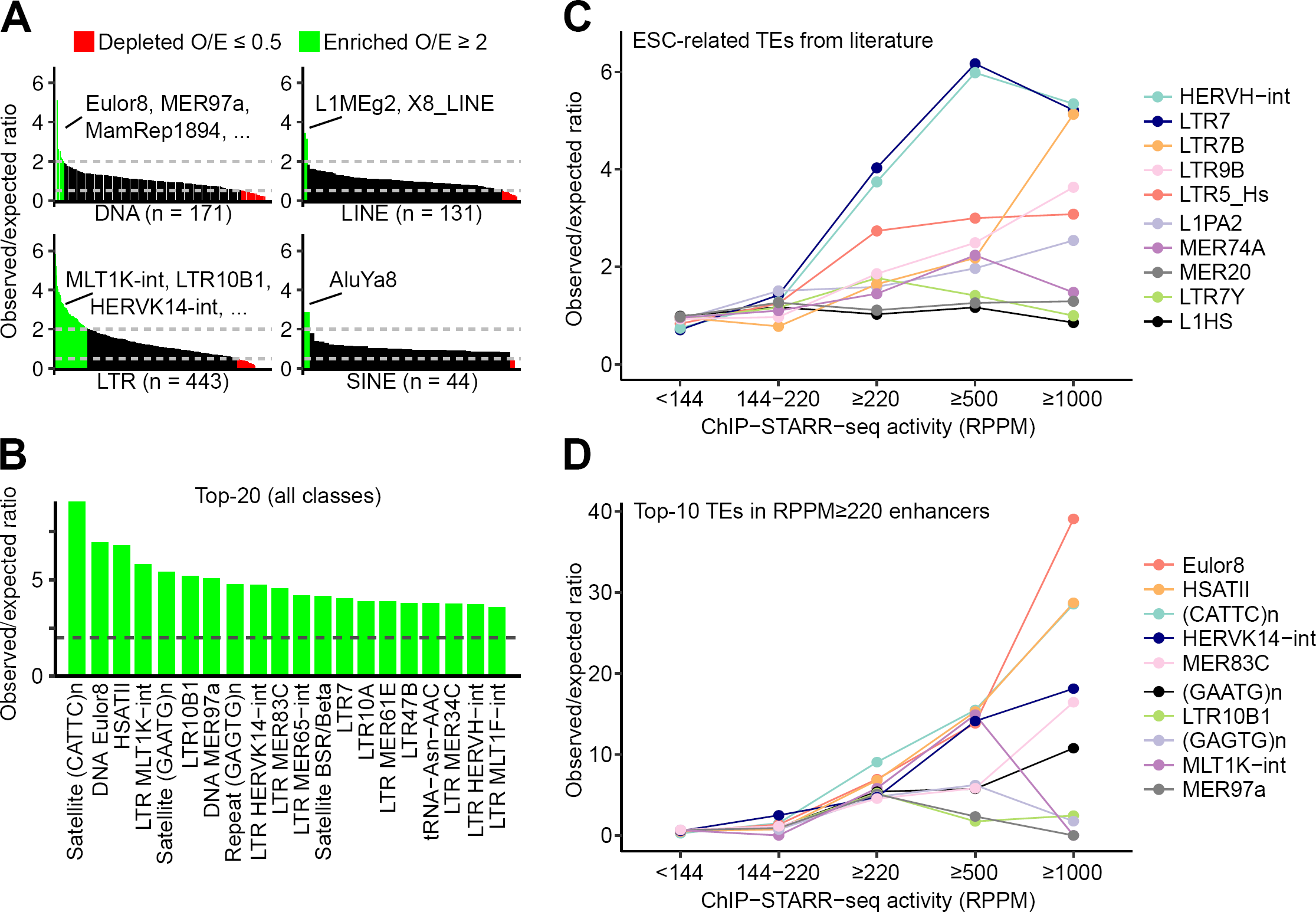
Distinct transposable elements are associated with enhancers of differing activity in hESCs. **A**) Bar graph showing enrichment (observed over expected; O/E) ratio for the occurrence of distinct transposable element (TE) families (LTR, DNA, SINE and LINE) in high activity ChIP-STARR-seq enhancers (RPPM ≥220). TEs with a high O/E (≥2; green) or low O/E (≤0.5; red) are shown. **B**) Bar graph of the top-25 most enriched TE families in high activity enhancers (RPPM ≥220). **C**)Line graph of the enrichment ratio for distinct transposable element families in enhancers with various activity levels. **D**) As panel C but for the top-10 most enriched families of transposable elements in B.

### ChIP-STARR-seq dissects super-enhancers into small functional units

Recently, large linear tracts of chromatin, referred to as super-enhancers (SEs) have been identified that function to regulate lineage-specific gene expression (Hnisz et al., 2013; Whyte et al., 2013). Compared to traditional enhancers, SEs have increased binding of Mediator, specific histone marks and lineage-specific TFs. Whether the full length of SEs is required for biological activity has become a matter of debate (Dukler and Gulko, 2016; Hay and Hughes, 2016; Moorthy et al., 2017; Shin et al., 2016). As a further application of our enhancer catalogue, we used the data to dissect the regulatory potential of DNA underlying SE regions. SEs were first identified by H3K27ac enrichment in primed (Figure 6A, **File S1**) and naive (Figure S8A) hESCs. Alignment of ChIP-STARR-seq data to these SEs showed that the H3K27ac intensity used to define SEs correlated to RPPM levels (Figures 6B, S8B), supporting the notion that SE-likeness is an indicator of functional potential. However, as exemplified by the SE covering the *FGFR1* gene, detailed examination indicates that strong RPPM signals originate from only a small region within the entire SE (Figure 6C). Therefore, luciferase assays were used to determine the enhancer activity of DNA in the neighborhood of this active region. Strong activity was confined to a 596bp region with other DNA elements from this SE devoid of enhancer activity (Figure 6D). This indicates that the *FGFR1* SE is composed of small units with enhancer activity. To test whether this finding is valid globally, the relative abundance of highly active plasmids (RPPM ≥220) in SEs compared to “normal” enhancers (NEs) was examined. Most enhancers contained only a small percentage of active plasmids within their bounds (Figures 6E). Although this fraction was slightly higher in SEs than in NEs, it accounted for only a minority (∼3%) of the genome annotated as SEs. Therefore, only a small part of the large SEs has enhancer function (Figure 6F). These conclusions also apply to naive hESC SEs (**Figure S8**). Although H3K27ac intensities in naive hESCs were, in general, slightly less than in primed hESCs (Figure S8C), the majority of SEs (n=2,597) were found in both cell states (Figure S8D). As for primed hESCs, only a minor portion of naive hESC SEs possessed high enhancer activity (mean across SEs =1.9%; (Figure S8E).

**Figure 6:**
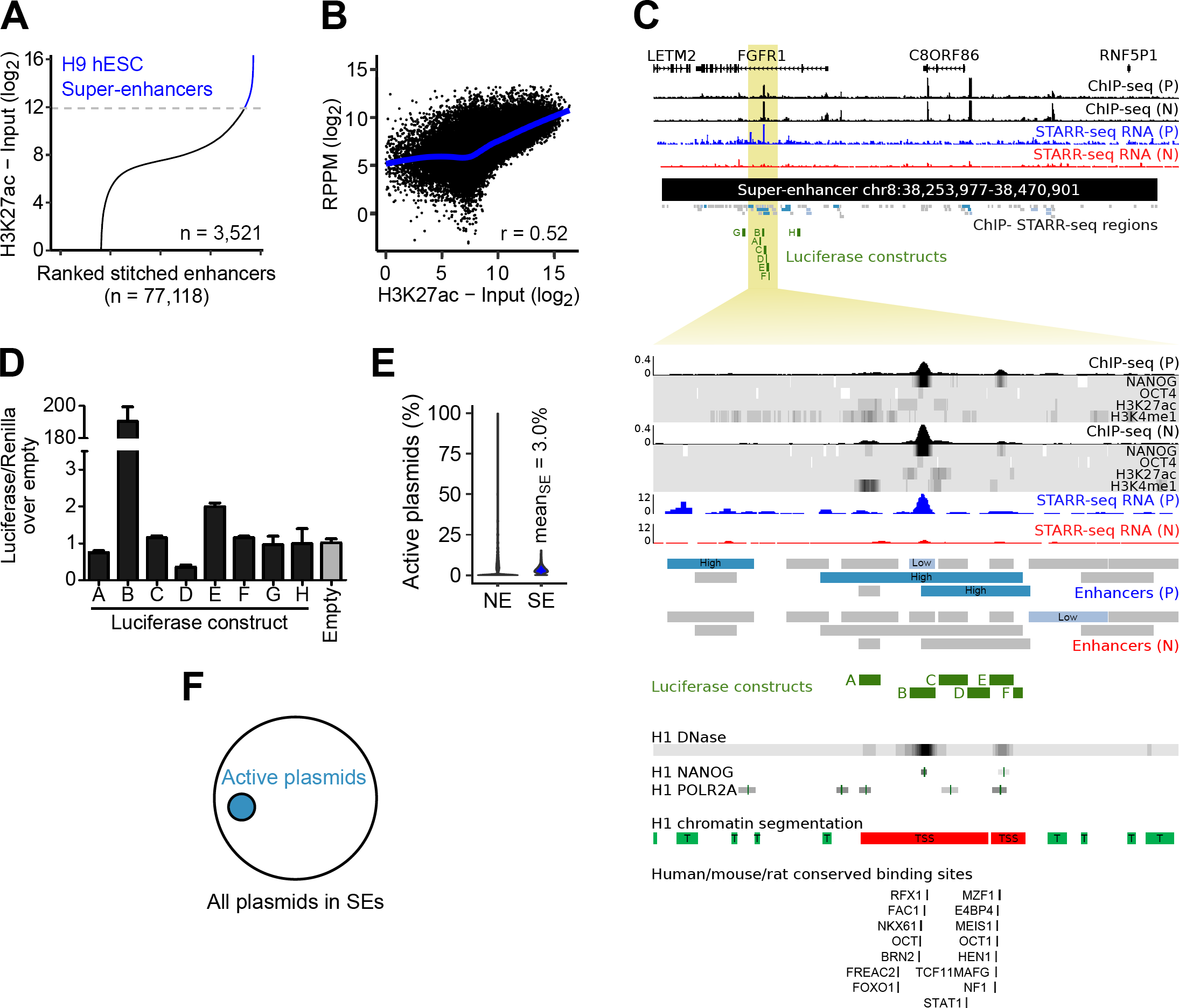
ChIP-STARR-seq dissects super-enhancers into functional elements. **A**)Super-enhancers (SEs) were called from H3K27ac ChIP-seq data using ROSE (Whyte et al., 2013). **B**)Scatterplot contrasting SE intensity (H3K27ac enrichment over input) with ChIP-STARR-seq activity. The Pearson correlation (*r*) is shown with the blue line indicating a generalised additive model fit to the data. **C**)Genome browser view of the FGFR1 super-enhancer, with ChIP-seq tracks for NANOG, OCT4, H3K27ac, and H3K4me1 in primed and naive hESCs. The top plot shows the SE locus, the bottom plot zooms into the indicated region within the second intron of FGFR1. Also shown are the positions of all regions assessed by ChIP-STARR-seq (grey) and active enhancers (blue) from this study and the coordinates of eight luciferase constructs matching selected enhancers (labeled A-H). Also included are public data available via the UCSC genome browser (from top to bottom): H1 DNase hypersensitivity, ChIP-seq for NANOG and POLR2A, ENCODE chromatin segmentation and DNA sequence motif occurrences. Enhancer activities are concentrated at a single position within the FGFR1 super-enhancer. **D**) Luciferase assays of DNA sequences depicted in green in C). Enrichment in luciferase activity relative to empty vector, normalized to the Renilla transfection control. N=2, error bars represent SD. **E**) Violin plots showing the proportion of active plasmids (RPPM ≥220) for 3,521 super-enhancers (SE) compared to normal enhancers (NE). Each data point is one SE or NE region. **F**)Venn diagram of the active subspace (covered by plasmids with RPPM ≥220) of the entire SE space (all plasmids occurring within SEs).

## Discussion

We present here a large-scale analysis of enhancer activity in human embryonic stem cells. Previous studies to elucidate the *cis*-regulatory network of hESCs used correlations between the presence of histone modifications, to predict potential enhancers (Hawkins et al., 2011; Rada-Iglesias et al., 2011; Xie et al., 2013). Only a small proportion of predicted enhancers sequences were functionally validated (Attanasio et al., 2013; Rada-Iglesias et al., 2011; Visel et al., 2007). Therefore, the degree to which correlations with histone marks can predict enhancer function genome-wide has remained unclear. Indeed, sequences not marked by H3K4me1 and H3K27ac can act as transcriptional enhancers (Pradeepa et al., 2016). Here we have combined ChIP with STARR-seq as a direct test of the ability of DNA sequences bound by OCT4, NANOG or by the histone marks H3K4me1 and H3K27ac to function as enhancers. The glucocorticoid receptor network in lung epithelial cells was recently assessed similarly (Vockley et al., 2016). Importantly, we found that only a subset of these sequences displayed enhancer activity. We find that TF binding is closely linked with enhancer activity, in line with recent reports (Kwasnieski et al., 2014) (Dickel et al., 2014; Ernst et al., 2016; Kheradpour et al., 2013) (Vockley et al., 2016). However, assigning enhancer potential based on the presence of histone marks or TF binding alone identifies putative enhancers that cannot enhance transcription when tested functionally. In addition, previous approaches did not identify some enhancer classes, as illustrated by our discovery of a previously unrecognized group of functional enhancers associated with housekeeping genes. The HK module is characterized by reduced binding of pluripotency-associated TFs and histone marks. This reduced binding likely placed these regions below the detection threshold in previous ChIP-seq studies that lacked a functional read-out.

Previous studies identified crucial roles for OCT4, NANOG and SMAD3, the latter of which are downstream mediators of TGF-β signaling in the maintenance of hESC pluripotency (James et al., 2005; Mullen et al., 2011; Xu et al., 2008). High enhancer activity is enriched near the binding peaks of these TFs, suggesting that they contribute directly to enhancer function. Other MPRA studies have shown that heterotypic clusters of different TF binding sites can drive higher enhancer activity (Smith et al., 2013) and it will be of future interest to decipher the contributions of individual TF binding to these active enhancers.

Several classes of TEs were also enriched at active enhancers, as reported recently (Ernst et al., 2016). TEs are enriched in species-specific TF binding sites and have been hypothesized to shape the enhancer network in hESCs (Glinsky, 2015; Kunarso et al., 2010). Our data indicate that only a limited number of TEs contribute to enhancer function providing the means to further refine the rewiring hypothesis.

A further way in which our findings identify additional features of enhancers is in the position of enhancers relative to transcription units. Most enhancers studied to date lie within distal elements or intronic sequences. However, some sequences near TSSs are detected by the ChIP-STARR-seq assay. As test enhancers are inserted downstream of the GFP ORF in the STARR-seq plasmid (Figure 1A) GFP-positive transcripts cannot be made by initiating transcription *in situ* from the inserted TSS. Therefore, sequences near a TSS can exert enhancer activity, in line with a recent report (Engreitz et al., 2016). Furthermore, a subset of housekeeping enhancers lie close to a TSS, suggesting that nearby enhancers may regulate some human housekeeping genes. It will be interesting to determine whether the specific links identified between enhancers and core-promoters that distinguish housekeeping genes from developmental genes in *Drosophila* also exist in mammalian cells (Cubenas-Potts et al., 2016; Zabidi et al., 2015).

Several groups have recently developed cultures supporting a more naive hESC state (Gafni et al., 2013; Takashima et al., 2014; Theunissen et al., 2014) with cells cultured in some of these conditions able to contribute to interspecies chimaeras (Gafni et al., 2013; Wu et al., 2017). Here we have used one such culture condition to compare enhancer activity in primed and naive cells. Enhancer activity alters substantially between primed and naïve hESCs. Pluripotency in both states is established by differential use of regulatory elements that is partly reflected in gene expression changes. Many active enhancers in naive cells are located close to TSSs, which may relate to the reported decrease in bivalent marks near TSSs in hESCs cultured in NHSM (Gafni et al., 2013). Further studies should clarify differences between different states of naive pluripotency and how these relate to differences in enhancer usage.

SEs are characterized by large domains marked by H3K27ac with increased binding of Mediator and other TFs. ChIP-STARR-seq shows that the majority of sequences covered by SEs lack enhancer activity. Rather, enhancer activity is limited to small domains within the SEs that frequently overlap with TF binding sites. This suggests that the observed chromatin signatures at SEs might be a consequence of enhancer activity from much smaller units. Recent reports suggest that SE constituents may function alternatively as either independent and additive enhancers (Hay and Hughes, 2016; Moorthy et al., 2017), as constituents in a temporal and functional enhancer hierarchy (Shin et al., 2016), or as interdependent units (Hnisz et al., 2015) exhibiting synergy (Suzuki et al., 2017). The large scale identification of such small active constituents within SEs reported here will be a valuable resource to further decipher the regulatory mechanisms contributing to SE formation, evolution and function.

The catalogue of functional enhancers presented here provides the means to refine models of the regulatory circuitry of hESCs and a framework for deepening understanding of transcriptional regulation in humans. Given the increasing appreciation of the importance of the regulatory genome in health and disease we expect that this resource and the more widespread use of MPRAs such as ChIP-STARR-seq will advance basic and translational research alike.

## Author contribution

TSB and IC conceived the study. TSB performed the molecular biology and contributed to the bioinformatics analysis and cell culture. AR developed the primary data analysis pipeline. FH performed bioinformatics analysis, visualization and interpretation. MZ performed cell culture, immunofluorescence and generated CRISPR/Cas9 deleted clones. CB supervised the bioinformatics analysis. TSB, FH and IC wrote the paper, with input from all authors.

## Acknowledgements

We thank S. Pollard, D. O’Carroll, A. Soufi and S. Tomlinson for comments on the manuscript, E. Hall-Ponsele and F. Rossi for technical support and R. Pantier, J. Zhang and other members of the Chambers’ lab for helpful discussions. We thank A. Stark (IMP) for the STARR-seq plasmid, F. Zhang (Broad Institute) for eSpCas9(1.1) and Y. Wang and D. Hay for H9 hESC cells. IC’s lab is supported by the Medical Research Council (UK) and The Wellcome Trust. TSB was supported by fellowships from Niels Stensen, EMBO (EMBO-LTF) and Marie Sklodowska-Curie (H2020 MSCA-IF). FH was supported by the DFG (grant HA 7723/1-1). CB is supported by a New Frontiers Group award of the Austrian Academy of Sciences and by an ERC Starting Grant (no. 679146). Sequencing was done by Edinburgh Genomics.

## Materials & Methods

### Cell Culture

H9 human embryonic stem cells were cultured on Matrigel coated cell culture plates, using mTesR1 medium (Stem Cell Technology, 05850). Cells were routinely split (ratio 1:3-1:4) using 0.5mM EDTA (Invitrogen, 15575020). For transfection, single cells were obtained by Accutase treatment (Invitrogen, A1110501), in the presence of Rock inhibitor, Y-27632 (10uM, Cambridge bioscience, SM02-10). For conversion to the naive state, cells were split on irradiated MEFs on gelatin coated plates and media was changed to NHSM media, as described by Gafni et al. (Gafni et al., 2013), containing knockout DMEM (Invitrogen), 20% knockout serum (Invitrogen), human insulin (Sigma, 12.5μg ml-1 final concentration), 20 ng ml-1 recombinant human LIF (Millipore), 8 ng ml-1 recombinant bFGF (Peprotech) and 1 ng ml-1 recombinant TGF-β1 (Peprotech), 1 mM glutamine (Invitrogen), 1% nonessential amino acids (Invitrogen), 0.1 mM beta-mercaptoethanol (Invitrogen), penicillin-streptomycin (Invitrogen) and small molecule inhibitors: PD0325901 (1μM, ERK1/2i, Axon Medchem); CHIR99021 (3μM, GSKβi, Axon Medchem); SP600125 (10μM, JNKi, Abcam ab120065) and SB203580 (10 μM, p38i,Abcam ab120638) Y-27632 (5μM, ROCKi) and protein kinase C inhibitor G06983 (5 μM, PKCi, Abcam, ab144414). Cells were 1:10 passaged using TrypLE™ (Invitrogen, 12604021) in the presence of Rock inhibitor and maintained for more than 10 passages in NHSM media prior to analysis. All cells were regularly karyotyped and checked for the presence of mycoplasm.

### Chromatin immunoprecipitation

For chromatin immunoprecipitation, 2x10^7 H9 primed or naive hESC were harvested in 9 ml of medium and cross-linked by addition of 270 μl 37% Formaldehyde (Sigma, final concentration of 1%), for 10 min at room temperature under rotation. 1 ml of 1.25 M Glycine was added, cells were incubated on ice for 5 min and 3x washed with ice cold PBS. At this point, cross-linked cell pellets were snap-frozen and stored at -80°C, or immediately processed for sonication. Prior to sonication, cells were resuspended in 1ml TE-I-NP40 (10mM TRIS-HCl pH 8, 1mM EDTA, 0.5% NP40, 1mM PMSF, 1x Protease inhibitor complex (PIC, Complete tablets, 04693116001, Roche)) incubated on ice for 5 min and centrifuged for 5 min at 2500 rpm at 4°C in a refrigerated bench top centrifuge (Eppendorf). Supernatant was removed and nuclei were resuspended in 1 ml ice-cold lysis buffer (50mM TRIS-HCl pH 8, 10mM EDTA, 1% SDS, 1mM PMSF, 1x PIC) and transferred to a 15 ml Falcon tube for sonication, using a Diagenode Bioruptor Next Gen (40 cycles of 30” on, 30” off). After transfer to an Eppendorf tube and centrifugation for 10 min at 13200 rpm at 4°C, chromatin solution was aliquoted and used for immunoprecipitation or snap-frozen and stored at -80°C. A 20 µl sample was taken and served as a total input control. For immunoprecipitation, Protein Dynabeads G (10004D, Life Techology) were washed with PBS and incubated for 6 hours with 5 μg of antibody, at 4°C on a rotating wheel. Antibodies used were: goat-anti-NANOG (AF1997, R&D Systems), rabbit-anti-OCT4 (AB19857, Abcam), rabbit-anti-H3K4me1 (AB8895, Abcam) and rabbit-anti-H3K27ac (AB4729, Abcam); as a control, respective IgG antibodies were used (rabbit-IgG: 10500C, Life Technology, goat-IgG: SC-2028, Santa Cruz Biotechnology). After washing with PBS, antibody-coupled beads were incubated with 200 μl chromatin solution, diluted to a final volume of 2 ml with dilution buffer (167mM NaCl, 16.7mM TRIS-HCl pH 8.1, 1.2mM EDTA, 0.01% SDS, 1.1% Triton-X100, 1mM PMSF, 1x PIC), overnight at 4°C on a rotating wheel. Washing of beads was performed by incubation with ice-cold 1 ml of washing buffer, for 5 min, at 4°C on a rotating wheel, followed by removal of supernatant using a magnetic stand, for each of the following: 2x with wash buffer 1 (10mM TRIS-HCl pH 7.6, 1mM EDTA, 0.1% SDS, 1% Triton-X100, 0.1% NaDeoxychloate), 2x with wash buffer 2 (10mM TRIS-HCl pH 7.6, 1mM EDTA, 0.1% SDS, 1% Triton-X100, 0.1% NaDeoxychloate, 150mM NaCl), 2x with wash buffer 3 (250mM LiCl, 0.5% NP40, 0.1% NaDeoxychloate), 1x with TE 1x with 0.2% TritonX-100 and 1x with TE 1x, after which beads were resuspended in 100ul TE1x. Immuno-precipitated chromatin and total input control were decross-linked, by addition of 3 μl of 10% SDS and 5 μl Proteinase K (20 μg/μl, Roche) and 10 μl RNAse A (50 μg/μl, Roche) to each tube and incubation overnight at 65°C on a shaking thermomixer block, 1400 rpm (Eppendorf). The next day, beads were briefly vortexed and supernatants were transferred to new tubes using the magnetic stand. 100μl of TE1x containing 500mM NaCl was added to the beads and briefly vortexed, after which the supernatant was added to the first fraction of collected supernatant. Following Phenol / chloroform extraction, DNA was precipitated using 1μl glycogen (20mg/ml), 1/10 vol NaOAc (3M) and 100% ice-cold Ethanol, at -20°C for 1 hour, followed by centrifugation at 13200 rpm for 1 hour at 4°C. After a final wash with 70% ethanol, the DNA pellet was dried and resuspended in 50μl H_2_O. Concentration of ChIP DNA was determined by Qubit measurement following manufacturer’s instructions and sonication was assessed by gel-electrophoresis of total input DNA (target fragment size between 200 and 600 bp).

### ChIP-qPCR

Concentration of ChIP and total input control DNA was assessed by Qubit measurement (Life-Tech) according to manufacturer’s instructions and was diluted to 2 ng/μl. 2 μl of DNA was used per qPCR reaction, using a 2x Takyon qPCR master mix (No ROX SYBR, UF-NSMTB0701, Takyon). qPCR reactions were run on a Roche Lightcycler 480 II (Roche), using the following cycle conditions: 95°C 3 min, (95°C 10 sec, 60°C 30 sec, 72°C 25 sec) x45, followed by a melting curve from 95° to 65°C. All data shown are averages of at least 2 biological replicates and 3 technical replicates. All primers used are shown in **Table S5.**

### ChIP-seq library and ChIP-STARR-seq plasmid library preparation

For ChIP-seq and ChIP-STARR-seq plasmid library generation, 10 ng of ChIP DNA was used as starting material. Using NEB Next ChIP-seq library preparation kit (E6200 or E6240, NEB), DNA was end-repaired, dA-tailed and adapter-ligated according to manufacturer’s instructions. After adapter ligation and purification using AMPure-XP beads (0.8x, Beckman Coulter) and elution into 30μl of 0.1xTE, 25 μl of the reaction product was used for ChIP-seq library preparation, by PCR amplification with Illumina index primers (7335 and 7500, NEB) using the NEB Next Q Hot start high fidelity master mix (M0543S, NEB) according to manufactures instructions (cycle conditions: 98°C 30 sec, (98°C 10 sec, 65°C 75 sec) x15, 65°C 5 min, 4°C hold). After an additional round of AMPureXP bead purification, DNA was eluted in 0.1xTE without further size selection. Quality and quantity of the prepared ChIP-seq libraries was assessed on an Agilent Tapestation. All sequencing occurred on an Illumina HiSeq 2500 platform, using 50bp single-end sequencing.

The remaining 5 μl of purified adapter ligated DNA were used for ChIP-STARR-seq plasmid library generation. Therefore, DNA was diluted to a total volume of 10 μl in 0.1xTE and used as an input in 8 x 50μl PCR reactions using Phusion Polymerase, High-fidelity buffer (M0530L, NEB) and primers 147 STARRseq libr FW (TAGAGCATGCACCGGACACTCTTTCCCTACACGACGCTCTTCCGATCT) and 148 STARRseq libr RV (GGCCGAATTCGTCGAGTGACTGGAGTTCAGACGTGTGCTCTTCCGATCT) (Arnold et al., 2013), which prime on the adapter sequences and add a 5’and 3’ 15 nucleotide homology sequence to the reaction products which are used for Gibson assembly. After PCR amplification (cycle conditions: 98° 2 min, (98°C 10 sec, 62°C 30 sec, 72°C 30 sec) x 15, 72°C 5 min, 4°C hold), PCR reactions were pooled, purified using AMPure XP beads (1.8x), eluted in 30 μl 0.1xTE and used for Gibson assembly. Therefore, 15 μg of the mammalian STARRseq plasmid (a kind gift of A.Stark) (Arnold et al., 2013) were digested with AgeI-HF and SalI-HF (NEB) for 8h at 37°C, column purified (Nucleospin purification columns, 740609250, Machery-Nagel), eluted in 30 μl elution buffer and used as a vector in a Gibson reaction, using 2 μl of digested plasmid, 5 μl purified PCR product, 3 μl H20 and 10 μl of a home-made Gibson reaction (100mM Tris-HCl, 10mM MgCl2, 0.2 mM dNTP (each), 0.5U Phusion DNA polymerase (NEB), 0.16U 5’ T5 exonuclease (Epicentre), 2 Gibson reactions per library. After incubation at 50°C for 1 hour, Gibson reaction were pooled and precipitated by addition of 1 μl Glycogen (20 μg/μl, Roche, 1090139300), 5 μl NaOAc (3M) and 125 μl ice-cold 100% ethanol, incubation at -20°C for 1 hour and centrifugation for 1 hour at 13200 rpm at 4°C, followed by a final wash in 70% ethanol. After air drying, DNA pellet was dissolved in 10 μl water and used for electroporation into electrocompetent MegaX DH10β E.coli bacteria (Invitrogen), according to manufacturer’s instructions, using a Biorad pulser. A total of 5 electroporations per library were performed with each 2 μl of DNA. After recovery in 1 ml SOCS medium each, bacteria were grown for 1 hour at 37°C in a bacterial shaker in the absence of antibiotics. Then, bacteria were pooled together and 50 μl of a 1:100 and 1:10000 dilution was plated on Ampicillin containing Agar plates to enable estimation of the number of transformants after overnight growth at 37°C (Control electroporations with Mock-Gibson without addition of PCR product plated on Ampicillin, or digested STARRseq plasmid transformations on Ampicillin- and Ampicillin/Chloramphenicol-containing Agar plates were negative, confirming complete digestion of the STARR-seq plasmid and a functional Ccdb counter-selection in DH10βE.Coli). The remaining 5 ml of bacteria culture were incubated in a total volume of 2 liter of LB-media supplemented with Ampicillin and allowed to grow for 16 hours in a bacterial shaker at 37°C. Plasmid DNA was isolated using a Qiagen Maxiprep kit according to manufacturer’s instructions and eluted in 500 μl 10mM Tris-HCl, pH 7.4. Concentration was determined by Nanodrop measurement.

### Transfection of plasmid libraries

Primed and naive H9 hESCs were transfected using either Nucleofection (Lonza, VPH-5022), or using Lipofectamin3000 according to manufacturer’s instructions. For each transfection, six million cells were used and transfected with 8 μg of plasmid library DNA and 500 ng pmCherry-N1 plasmid (Clonetech) as transfection control. Cells were incubated in 10 cm dishes and 24h post-transfection, single cells were harvested and subjected to FACS. Non-transfected cells were used to set sorting gates, DAPI was used as a marker for dead cells. All percentages mentioned are relative to the fraction of DAPI-negative, single cells.

### Preparation of ChIP-STARR-seq RNA and DNA samples for sequencing

A minimum of 400,000 GFP-positive, sorted cells were used to isolate total RNA using Trizol (Thermo Fisher) according to manufacturer’s instructions. The mRNA fraction was captured using Oligo (dT)25 beads (61002, Life Technologies) and DNAseI treated (18068-015, Life Technologies), followed by reverse transcription using 2 μl SuperscriptIII (18080-044, Life Technologies) using a GFP-mRNA specific primer (149 STARRseq rep RNA cDNA synth, CAAACTCATCAATGTATCTTATCATG) at 50°C for 90 minutes, in a total reaction volume of 21 μl. To repress residual plasmid DNA contamination, cDNA was PCR amplified using a combination of primers (152 STARR reporter specific primer 2 fw, GGGCCAGCTGTTGGGGTG*T*C*C*A*C and 153 STARR reporter specific primer 2 rv, CTTATCATGTCTGCTCGA*A*G*C, where * represent phosphorothioate bonds) spanning a synthetic intron in the STARR-seq plasmid, as previously described (Arnold et al., 2013). PCR was performed with Phusion polymerase and High-fidelity buffer, in 6 x 50 μl reactions (cycling conditions: 98°C 2 min, (98°C 10 sec, 62°C 30 sec, 72°C 70 sec) x15, 72°C 5 min, 4°C hold). PCR reactions were pooled, purified using AMPureXP beads (1.0x) and eluted in 18 μl 0.1xTE. Absence of significant plasmid contamination in the PCR amplified cDNA was assessed by qPCR using a primer-set amplifying an amplicon from the STARR-seq plasmid backbone (161 STARRseq detect plasmid backbone qPCR fw, CATCATCGGGAATCGTTCTT, and 162 STARRSeq detect plasmid backbone qPCR rv, TGAAGATCAACTGGGTGCAA), relative to a primer-set amplifying GFP (154 STARRseq GFP fw, ACGGCCACAAGTTCTCTGTC, and 155 STARRseq GFP rv, GCAGTTTGCCAGTAGTGCAG). PCR amplified cDNA was then used in a second round of PCR to add Illumina index primers (7335, 7500, NEB) using priming on the adapter sequences added during the plasmid library generation. PCR was performed in 1-4x 50 μl reactions using Phusion polymerase and High-fidelity buffer (NEB)(cycling conditions: 98°C 2 min, (98°C 10 sec, 65°C 30 sec, 72°C 30 sec) x13, 72°C 5 min, 4°C hold), after which PCR reactions were pooled, purified using AMPureXP beads (1.0x) and eluted in 15 μl 0.1xTE. Corresponding plasmid libraries were similarly amplified in a nested PCR, using primers detecting the STARR-seq plasmid (160 STARR reporter specific primer for plasmid DNA fw, GGGCCAGCTGTTGGGGTG, and 153 STARR reporter specific primer 2 rv, CTTATCATGTCTGCTCGA*A*G*C, where * represent phosphorothioate bonds) and Illumina index primers. In addition to sequencing libraries prepared from plasmid maxiprep DNA, we also sequenced plasmid libraries reisolated from transfected hESCs. For this, we transfected H9 hESCs as described above and harvested non-sorted cells 24h post-transfection, followed by plasmid reisolation using a Qiagen miniprep isolation kit and sequencing library preparation. Quantity and quality of generated sequencing libraries was assessed on an Agilent Tapestation. All sequencing occurred on an Illumina HiSeq 2500 platform, using 50bp or 125 bp paired-end sequencing. Up to 22 RNA samples were pooled on a single lane. During data-processing all reads were trimmed to 50bp length to improve consistency.

### RT-qPCR

For RNA analysis of complete cultures, cells were lysed in Trizol (Thermo Fisher) and RNA was prepared according to manufacturer’s instructions. 1 μg of RNA was treated with DNAseI (Invitrogen) to remove genomic DNA contamination and cDNA was obtained through reverse transcription using SuperScriptIII (Invitrogen) in the presence of RNAseOUT (Invitrogen). cDNA was diluted in DEPC-treated water to a final volume of 200 μl and 2 μl of cDNA was used per qPCR reaction, using a 2x Takyon qPCR master mix (No ROX SYBR, UF-NSMTB0701, Takyon). qPCR reactions were run on a Roche Lightcycler 480 II (Roche), using the following cycle conditions: 95°C 3 min, (95°C 10 sec, 60°C 30 sec, 72°C 25 sec) x45, followed by a melting curve from 95° to 65°C. All data shown are averages of at least 2 biological replicates and 3 technical replicates, normalized to TBP. All primers used are shown in **Table S5.**

### Immunostaining

Cells were grown on culture dishes suitable for confocal microscopy (Ibidi, 81156) and fixed using 4% v/v Paraformaldehyde at room temperature for 10 min. After permeabilisation using 0.3% Triton/PBS and incubation with blocking solution (1% BSA, 3% Donkey serum, 0.1% triton in PBS), cells were incubated with primary antibody O/N at 4°C. After washing with PBS, cells were incubated with secondary antibody at RT for 1h, washed and counterstained with DAPI. Imaging occurred on a Leica SP8 STED-CW confocal microscope and images were processed using ImageJ software. Antibodies used are: goat-anti-NANOG (1: 200, AF1997, R&D Systems), rabbit-anti-OCT4 (1: 200, AB19857, Abcam). Secondary antibodies were Donkey-anti-goat conjugated to Alexa fluor488 (1:800, A11055, Invitrogen) and Donkey-anti-rabbit conjugated to Alexa fluor568 (1:1000, A10042, Invitrogen).

### Western blotting

Whole cell protein extracts were isolated and Western blotting was performed using standard procedures using pre-cast 10% Bis-Tris Bolt gels (Invitrogen). Primary antibody used was goat-anti-NANOG (1: 500, 1μg/ml, AF1997, R&D Systems), secondary antibody conjugated to fluorophores was donkey-anti-goat-IRDey680 (1:500, 926-68074, Li-cor). Rabbit-anti-Laminin B (1:1000, AB16048, Abcam) served as a loading control and was detected by chemiiluminescence. Imaging occurred on an Odyssey imager (Li-cor).

### Luciferase assays

Enhancer sequences were PCR amplified from human genomic DNA using Phusion polymerase and cloned by Gibson assembly into a KpnI-NheI linearized Pgl3 promoter luciferase vector. For primer sequences, see **Table S5.** All constructs were sequence-verified by Sanger sequencing and co-transfected with a Renilla expressing plasmid using Lipofectamin 3000 into H9 hESCs. 48h post-transfection illuminescence was assessed using the Dual Glo luciferase kit (E2920, Promega) according to manufacturer’s instructions, on a Promega Glumax Multidection system. All data shown are average from at least two biological replicates and two technical replicates, representing fold-change in luciferase activity compared to empty vector controls and normalized for Renilla transfection control.

### CRISPR/Cas9 genome editing

Oligonucleotides for gRNAs flanking the OCT4 distal enhancer (OCT4 DE human gRNA1 fw: CACCGGAGATGGGCACACGAACAG, OCT4 DE human gRNA1 rv: AAACCTGTTCGTGTGCCCATCTCC, OCT4 DE human gRNA2 fw: CACCGTCTGCGTCCCTCTCGGGAA, OCT4 DE human gRNA2 rv: AAACTTCCCGAGAGGGACGCAGAC) were annealed and cloned into a BbsI digested spCas9 plasmid, from which the gRNAs are separately expressed together with a eSpCas9(1.1)-t2a-mCherry or eSpCas9(1.1)-t2a-GFP (modified from Addgene plasmid #71814, (Slaymaker et al., 2016). All plasmids were sequence verified and 1 μg of each gRNA was used to transfect primed H9 hESCs in a 6-well plate using Lipofectamine 3000. 48h post-transfection, mCherry and GFP double positive cells were FAC sorted and cells were plated at low density in 10 cm dishes coated with Matrigel in conventional mTesR1 hESC medium. Emerging clones were expanded and genotyped by PCR using primers flanking the gRNA targets (440: OCT4 DE genotyping 2 fw, GGGTCAGTGGCTCTATCTGC; 441: OCT4 DE genotyping 2 rv, TTCAACCAAACAGCACCTCA) to detect the approximately 650 bp deletion of OCT4 DE enhancer. Candidate clones after PCR screening were Sanger sequenced and correct clones were expanded and used for conversion experiments to the naive hESC state.

### ChIP-seq and ChIP-STARR-seq plasmid and RNA data processing

We trimmed possible adapter contaminants from reads using Skewer (Jiang et al., 2014). Trimmed reads were then aligned to the GRCh37/hg19 assembly of the human genome using Bowtie2 (Langmead and Salzberg, 2012) with the “*‐‐very-sensitive*” parameter. For ChIP-STARR-seq plasmid and RNA libraries, only properly paired, concordantly aligning and uniquely mapping fragments were kept. Genome browser tracks were created with the *genomeCoverageBed* command in BEDTools (Quinlan and Hall, 2010) and normalized such that each value represents the read count per base pair per million uniquely mapped reads. Finally, the UCSC Genome Browser’s *bedGraphToBigWig* tool was used to produce a bigWig file.

### Definition of ChIP-STARR-seq enhancer and activity levels

For ChIP-seq and plasmid DNA-seq libraries, peak calling was performed with MACS2 (Zhang et al., 2008) with default parameters, using the respective input samples as background. In addition, we called peaks with MACS2 without the background samples (“*—nomodel ‐‐extsize 147*” parameters). For each peak set, we fixed the peak width to 500 bp from the peak summit for transcription factors and 1000 bp for histone modifications and removed peaks that overlapped blacklisted features as defined by the ENCODE project (Hoffman et al., 2013). ChIP-seq peaks are given in **File S1**.

To define a set of enhancers to compare in our analysis of ChIP-seq, plasmid DNA-seq and ChIP-STARR RNA-seq samples, we produced a set of peaks by merging (*i.e.* computing the union) of peaks for the same factor across cell types and experiment types (ChIP-seq and plasmid DNA-seq). Furthermore, we generated a set of ”false positive” peaks for each factor to be used as a background dataset in our subsequent analyses. We defined this dataset as those peaks that were called by MACS2 without the total input control and that did not overlap with the peaks called with the control sample.

We initially quantified the intensity of ChIP-seq, plasmid DNA-seq and ChIP-STARR RNAseq datasets in the enhancer peak regions by counting the number of aligned reads overlapping each enhancer region. To get a more accurate and precise measure of plasmid reporter intensity for further analysis, we then made use of our paired-end sequencing data to unequivocally link RNA-seq reads to the plasmid that they came from. To do so, we matched RNA-seq reads to plasmid reads with the exact same start coordinate of the first read and the exact same end coordinate of the second read. Comparing the counts for both made it possible to define a measure of RNA-seq activity relative to the abundance of plasmids in the library (reads per plasmid). To avoid distortion by differences in sequencing depth, we first scaled RNA-seq read counts by library size (RNA-seq reads per million (RPM), *R*) and plasmid counts by library size (plasmid RPM, *P*) to define our final measure of activity level as reads per plasmid million (*RPPM = R / P*). We then used the maximum observed RPPM value as an estimate of enhancer-peak-level activity. Since our individual replicate datasets were sparse, with the same plasmids infrequently measured in both replicates, but our overall coverage of enhancers was much better, we used RPPM from all datasets generated in the same cell type (so specific to either primed or naive H9 hESCs) for this purpose. We could do so because the ChIP-STARR-seq plasmid libraries are independent from the antibody target used to pull down the enriched DNA fragments, thus the plasmids in all libraries jointly report the activity of the same genome. To objectively define thresholds to distinguish highly active, lowly active and inactive genome regions, we made use of “false positive” background peaks defined earlier (from the MACS2 software). These are peaks usually removed from the analysis in other publications, because they display peak-like coverage not only in the DNA enriched for ChIP-seq targets but also in unenriched sonicated DNA (input control). The reasoning is generally that these regions represent DNA that is particularly well accessible, however, is not of genuine regulatory importance. We make the assumption that these peaks, even if they may contain regulatory enhancers, were slightly less likely to contain genuine functional enhancers than “true positive” peaks (that is, peaks in ChIP-seq dataset without concordant peak in the input control). We therefore fitted a log-normal statistical model to the distribution of RPPM values in the false positive peaks and determined thresholds at which 5% (p < 0.05) or 10% (p < 0.1) of these peaks would have been called “active”. We did this separately in primed and naive hESCs because the distributions of RPPM values were different, and we then used the threshold determined (primed: θ_high_ = 220, θ_low_=144; naive: θ_high_=96, θ_low_=62) throughout our analysis (see Figures 2A, S6D). The coordinates of all genome regions assessed with activity calls are given in **File S1**.

### Motif enrichment analysis for generated ChIP-seq data sets

BED files of ChIP-seq data sets were generated with 500 bp sequences centered on the narrow ChIP-seq peak, and used for motif enrichment analysis using CentriMo (http://memesuite.org/)(Bailey and Machanick, 2012), using default settings.

### Assignment of enhancers to genes

We used GREAT, version 3.0.0 (McLean et al., 2010) to assign regulatory elements identified in ChIP-STARR-seq to their putative target genes, using the following settings: basal plus extension, proximal 5kb upstream and 1kb downstream, plus distal up to 100kb. Publically available, processed RNA-seq data from primed human ESCs were downloaded (Gifford et al., 2013; Ji et al., 2016; Takashima et al., 2014) and their RPKM value distribution was plotted for the various ChIP-STARR-seq regions grouped by activity in RPPM. For naive hESCs, we used publically available microarray data from the original study describing gene expression in naive cells cultured under NHSM conditions (Gafni et al., 2013).

### Comparison to previously published ESC enhancers

The coordinates of putative enhancers were obtained from the supplementary data of Hawkins et al, Rada-Iglesias et al and Xi et al (Hawkins et al., 2011; Rada-Iglesias et al., 2011; Xie et al., 2013), and when necessary converted to the hg19 version of the human genome using the liftOver tool. Overlapping enhancers were merged into 76,666 putative enhancers and joint to our ChIP-STARR-seq enhancers using GenomicRanges (Lawrence et al., 2013) in R (see Figure S4A,B, Table S3). We refer to those enhancers that overlapped with previously published enhancers and showed a ChIP-STARR-seq activity of RPPM>=220 as the ESC enhancer module (n=7,596). Conversely, we refer to active enhancers (RPPM >= 220) that did not overlap with the previously published enhancers as the housekeeping (HK) enhancer module (n = 13,604).

### Functional enrichment analysis

To help understand the function and relevance of different groups of enhancers, we used three types of functional enrichment analysis (**Table S2**).

(a) We used LOLA (Sheffield and Bock, 2016) to determine the relative over-representation of ChIP-seq peaks related transcription factor binding and other elements of known regulatory function. To this end, we used the *codex*, *encode_tfbs*, and *encode_segmentation* databases contained in the LOLA Core database and tested for the enrichment of overlap in genome regions with a specific level of activity (high, low or inactive) over the background of all ChIP-STARR-seq peaks.

(b) We also used the Enrichr web interface (February 2017 version) (Chen et al., 2013) to test genes linked to enhancers of interest for significant enrichment in numerous functional categories. In all plots, we report the “combined score” calculated by Enrichr, which is a product of the significance estimate and the magnitude of enrichment (combined score *c = log(p) * z*, where *p* is the Fisher’s exact test p-value and *z* is the z-score deviation from the expected rank).

(c) We additionally used the GREAT web interface (version 3.0.0) (McLean et al., 2010) for gene ontology analysis, using the following settings: basal plus extension, proximal 5kb upstream and 1kb downstream, plus distal up to 100kb, including curated regulatory domains, and whole genome (hg19) as background.

### Motif over-representation analysis

To find motifs that occurred more frequently in one enhancer group than in the others, we used FIMO (v4.10.2) (Grant et al., 2011) to scan the DNA sequences of all ChIP-STARR-seq enhancers for occurrences of known DNA motifs from the HOCOMOCO database (v10) (Kulakovskiy et al., 2016) using default parameters. We than compared the count of motif hits at a significance threshold of p<=0.05).

### Enrichment analysis for transposable elements

The UCSC RepeatMask (hg19) was downloaded from the UCSC Table Browser, imported into Galaxy (usegalaxy.org) (Afgan et al., 2016) and joined to the ChIP-STARR-seq activity calls for primed hESCs. The frequency of the various repeat sequences was counted for either all ChIP-STARR-seq regions that could be measured, or for the various subgroups binned according to their activity. To calculate the expected number of repeats present in each of the various activity groups, we divided the observed repeat counts in the total group by the number of regions that could be measured in all ChIP-STARR-seq regions, and multiplied this for the number of regions present in each activity subgroup (**Table S4**). We then calculated the ratio between observed and expected (O/E), and considered repeats with O/E<0.5 as depleted, or O/E>2 as enriched. For the subsequent data interpretation, we only focused on transposable elements that were >15 times present in all ChIP-STARR-seq regions.

### Super-enhancer analysis

To call super-enhancers in primed and naive H9 hESCs, we used the ROSE software (v0.1) (Whyte et al., 2013) to combine (“stitch”) ChIP-STARR-seq enhancers within 12.5 kb of each other and excluding 2.5 kb around known transcription start sites. We then asked the software to quantify the ratio of the H3K27ac ChIP-seq signal in primed and naive hESCs over the total input control and to call super-enhancers. The coordinates of all stitched enhancers, as well as primed and naive super-enhancers are given in **File S1.**

### Statistics for qPCR and luciferase assays

qPCR and luciferase assay figures were plotted and statistics were calculated using GraphPad Prism 5 software, p<0.05 was considered significant.

### Data availability

High-throughput sequencing data generated in this study have been submitted to the Gene Expression Omnibus (GEO) under accession code GSE99631. Additional data, genome browser tracks and an interactive search tool for active enhancers in the proximity of genes are available from a supplementary website under the following URL: http://hesc-enhancers.computational-epigenetics.org

## Supplemental Figure legends

**Figure S1, related to Figure 1:**
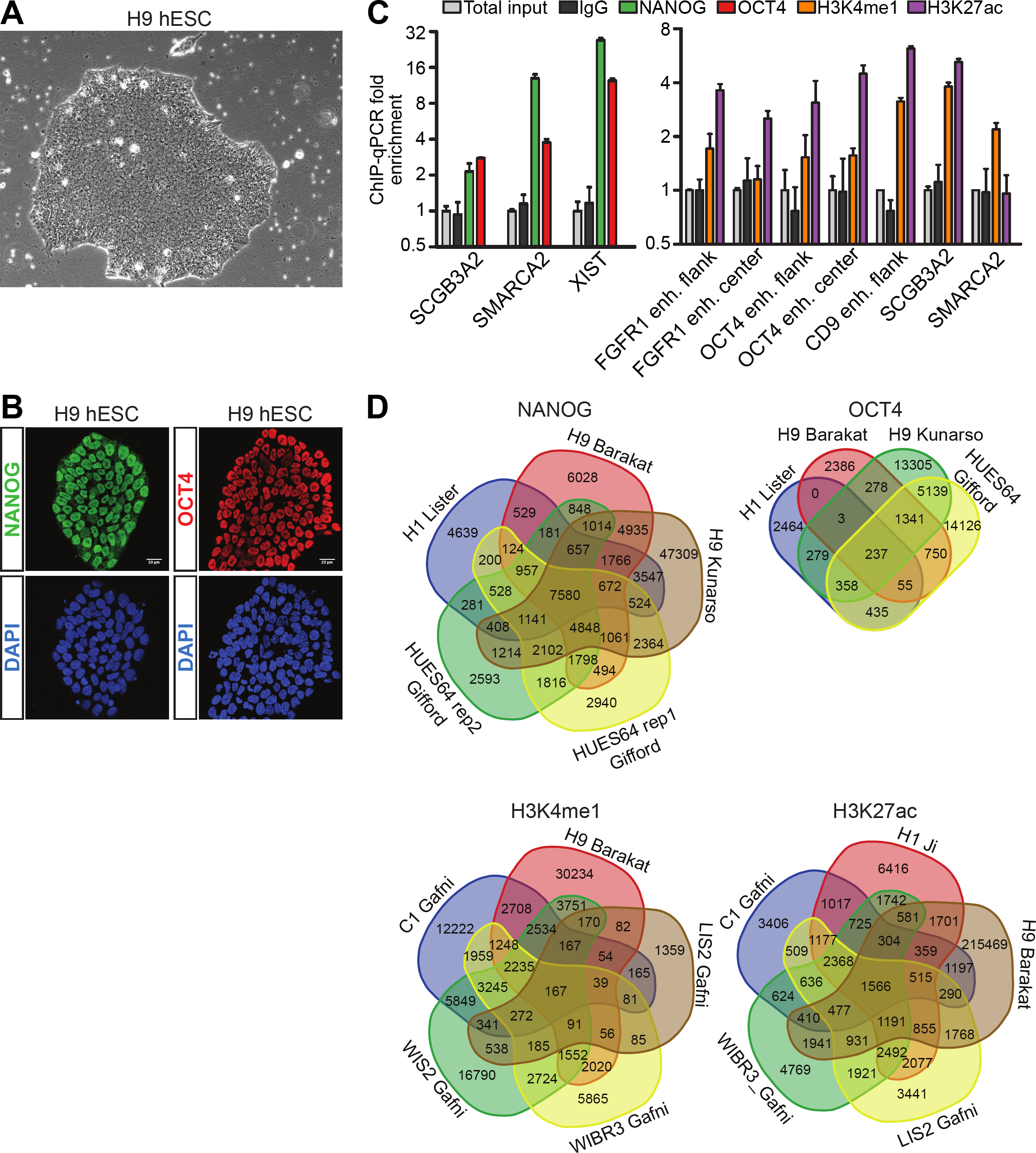
ChIP-seq in primed H9 human embryonic stem cells. **A**) Brightfield microscopy of a representative colony of H9 hESCs cultured on Matrigel in standard hESC culture conditions. **B**)Immunofluorescence of primed H9 hESCs for NANOG (green) or OCT4 (red); DNA is stained with DAPI (blue). **C**) ChIP-qPCR in primed H9 hESC with anti-NANOG, anti-OCT4 or rabbit IgG at known OCT4 and NANOG binding sites in *SCGB3A2, SMARCA* (Kunarso et al., 2010) and *XIST* (left) or with anti-H3K4me1, anti-H3K27ac or rabbit IgG at binding sites near *FGFR1* (at central and flanking locations), *POU5F1* (at central and flanking locations), *CD9*, *SCGB3A2*, and *SMARCA* (Rada-Iglesias et al., 2011) (right). The mean fold-enrichment is shown relative to total input control DNA, normalized to a non-bound site in *ACTB* (for OCT4 and NANOG), or *NCAPD2* (for H3K4me1 and H3K27ac). Error bars indicate standard deviations; n= 3. **D**) Venn diagrams of the overlap between ChIP-seq peaks in primed hESCs, indicating the cell line and study for NANOG, OCT4, H3K4me1 and H3K27ac. The numbers indicate overlapping peaks.

**Figure S2, related to Figure 1:**
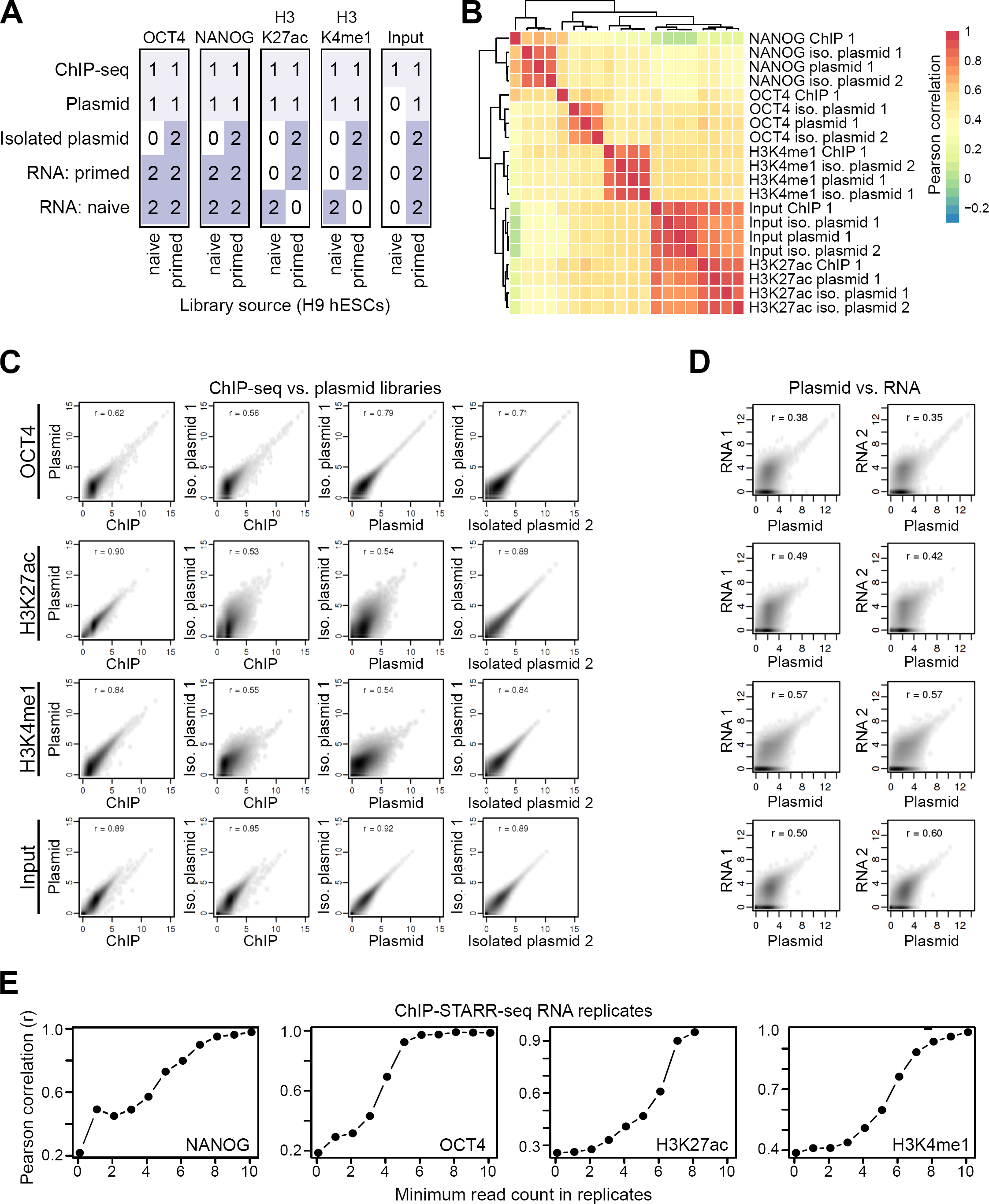
Overview of generated datasets. **A**) Summary table of the high-throughput sequencing datasets generated in this study, indicating the number of ChIP-seq, plasmid (corresponding to ChIP-STARR-seq plasmid libraries prior to transfection) and isolated plasmid (plasmid libraries 24h after transfection) samples, as well as RNA-seq data from GFP-positive primed and/or naive hESCs (RNA: primed and RNA: naive, respectively). **B**) Pearson correlation matrix of samples from the indicated ChIP-seq, plasmid libraries and isolated plasmid libraries from primed hESCs. Rows and columns have been arranged by hierarchical clustering with complete linkage. **C-D**) Scatterplots contrasting normalized read counts (reads per million) per peak for different datasets. The Pearson correlation coefficient (*r*) for each comparison is indicated for each plot. **C**) Comparison between ChIP-seq, DNA-seq of ChIP-STARR-seq plasmid libraries and two replicates of DNA-seq for isolated plasmid libraries post transfection, for OCT4, NANOG, H3K37ac, H3K27ac and genomic DNA (input). **D**) Comparison between ChIP-STARR-seq plasmid libraries prior to transfection and the corresponding RNA-seq read counts generated from two replicates of GFP-positive cells after transfection. **E**) Pearson correlation coefficients (*r*) between replicate STARR-RNA-seq measurements from the same pool of hESCs transfected with the same ChIP-STARR-seq library, shown as a function of minimum read count in both replicates (0 = unfiltered, 1 = at least one read from the same plasmid measured in each replicate, etc.).

**Figure S3, related to Figure 2:**
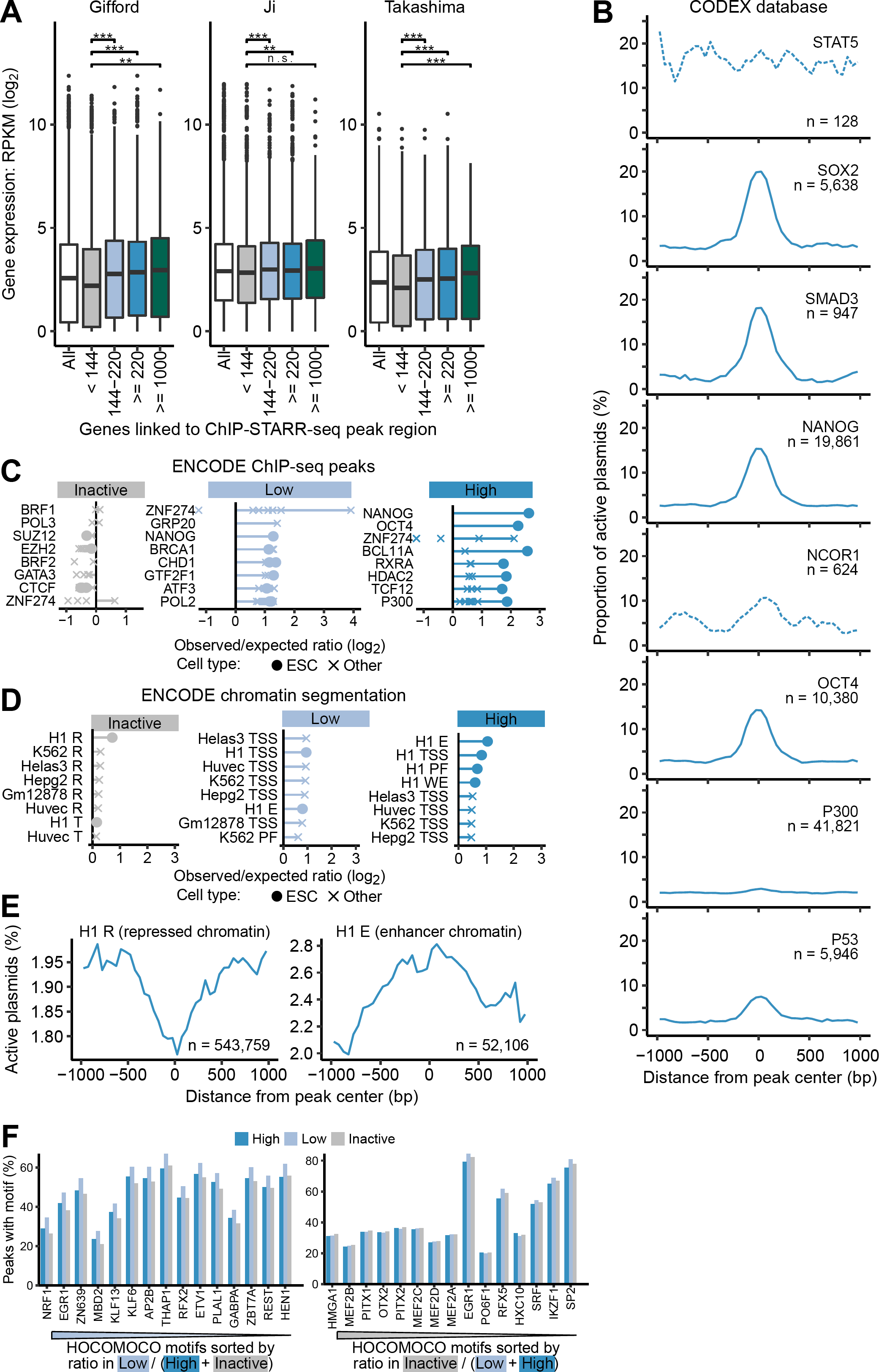
ChIP-STARR-seq in primed H9 hESCs. (Zhang et al., 2008) **A**) Boxplots showing the distribution of RNA-seq RPKM values of genes associated with enhancers grouped by activity level. Boxplots represent the interquartile range (IQR), the line is the median, whiskers extend to 1.5xIQR and outliers are indicated as dots. The numbers on the x-axis indicate thresholds on the RPPM activity level; RPKM, reads per kilobase million. RNA-seq datasets were from the following studies: HUES64 (Gifford et al., 2013); H1 (Ji et al., 2016); H9 (Takashima et al., 2014). **B**) Smooth line plots showing the proportion of active plasmids (RPPM ≥220) of all plasmids measured at the indicated distance from the peak center for ChIP-seq binding sites of factors from the CODEX database (Sanchez-Castillo et al., 2015) found preferentially associated at highly active enhancers (compare to Fig. 2F), averaged across all binding sites for the respective factor. The number of peaks (*n*) is indicated in each plot. **C**)Relative enrichment of DNA-binding proteins (DBPs) from the ENCODE database (2012) in inactive genome regions, as well as lowly active and highly active enhancers (compare to Fig. 2A). Shown are the log_2_-odds ratios between observed percentages of enhancers overlapping binding sites of each given DBP in the respective groups over the percentage of overlaps in the entire enhancer dataset. Each dot represents one ChIP-seq dataset for the given DBP and the lines connect the most extreme dot with zero for visualization. For each category, the eight most enriched DBPs are shown ranked by their mean log-odds ratio. ChIP-seq datasets produced from hESCs are indicated as dots and those from other cell sources as crosses. Enrichments were calculated using LOLA (Sheffield and Bock, 2016). **D**) LOLA enrichment plots as in panel C, but showing instead the relative over-presentation of ENCODE chromatin segments from different cell lines in enhancers with different activity levels. E, enhancer; PF, promoter-flanking region; R, repressed; T, transcribed; TSS, transcription start site; WE, weak enhancer. **E**) Line plots as in panel B, showing the proportion of active plasmids in a window around the center of repressed chromatin segments and enhancer chromatin segments from the ENCODE H1 chromatin segmentation (Hoffman et al., 2013). **F**) Top motifs from the HOCOMOCO database (Kulakovskiy et al., 2016) ranked by the frequency of hits within the lowly active enhancers compared to highly active enhancers and inactive genome regions (left) or within the inactive genome regions compared to lowly and highly active enhancers (right). The numbers indicate the proportion of peaks with at least one motif hit. Motif matches were determined using FIMO (Grant et al., 2011).

**Figure S4, related to Figure 3:**
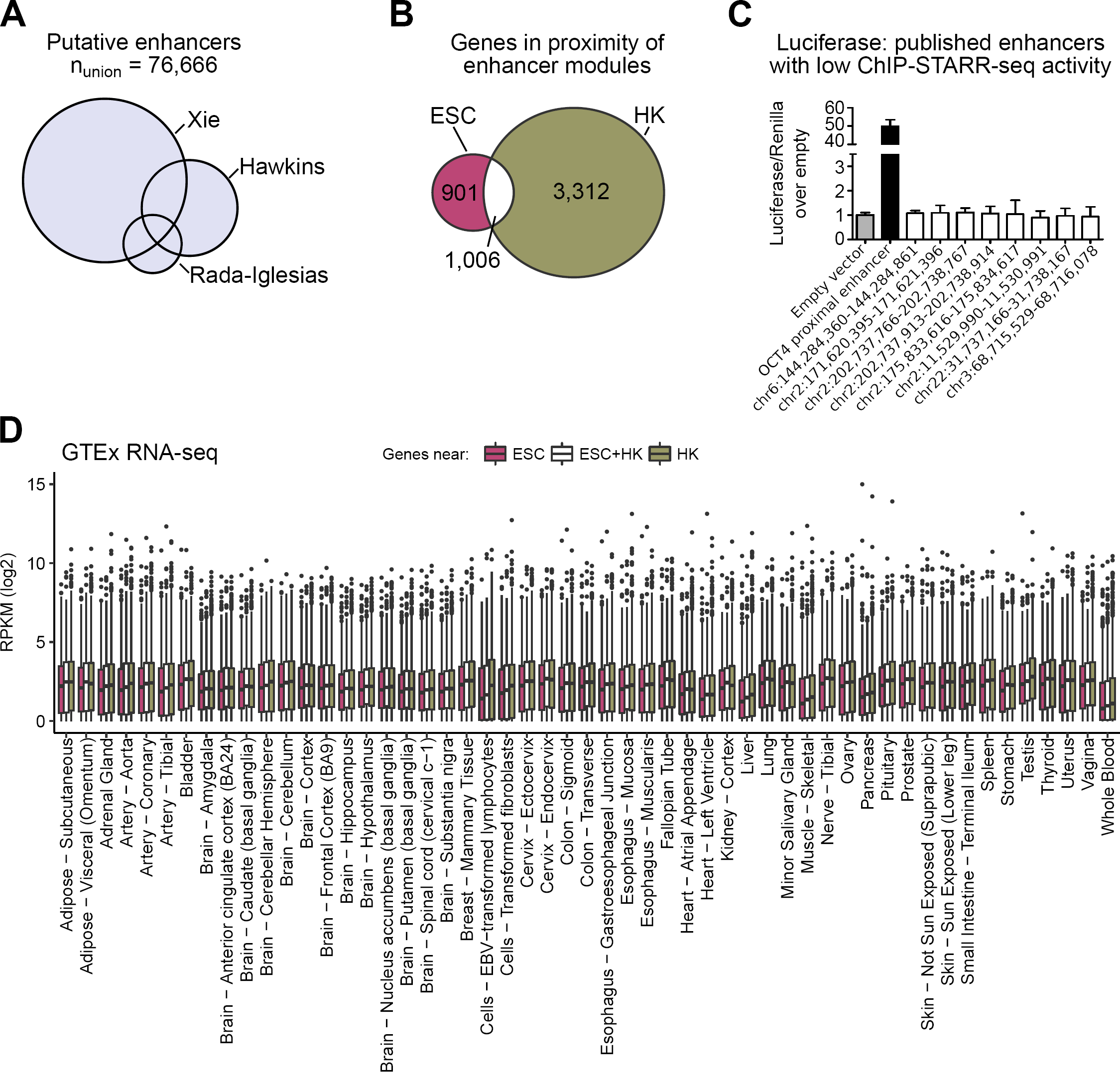
ESC-specific and housekeeping enhancer module. **A**) Illustration showing three source datasets (Hawkins et al., 2011; Rada-Iglesias et al., 2011; Xie et al., 2013) that contributed to the catalogue of putative ESC enhancers used in this study. We converted all enhancer coordinates to the same assembly (hg19) and then merged overlapping peaks resulting in a list of 76,666 putative enhancers. **B**) Illustration of the number of genes found in the proximity of ESC module enhancers (pink, left circle), HK module enhancers (olive, right), or with enhancers of both types (white, overlap). Enhancer-gene assignments were performed with GREAT (McLean et al., 2010). **C**) Luciferase assay in primed hESCs for eight putative enhancers that did not show activity in ChIP-STARR-seq. The OCT4 proximal enhancer (PE) is tested as a positive control. Luciferase activity is reported as the fold enrichment in luciferase counts over empty vector, normalised to the Renilla transfection control. Error bars indicate standard deviations, n=2. **D**) Boxplots of gene expression (RNA-Seq; log_2_) in different tissues from the GTEx database (2013) for all genes linked to ESC (pink) or HK (olive) module enhancers or both (white). Box-plots represent the interquartile range (IQR), the line is the median, whiskers extend to 1.5xIQR and outliers are indicated as dots. RPKM, reads per kilobase million.

**Figure S5, related to Figure 4:**
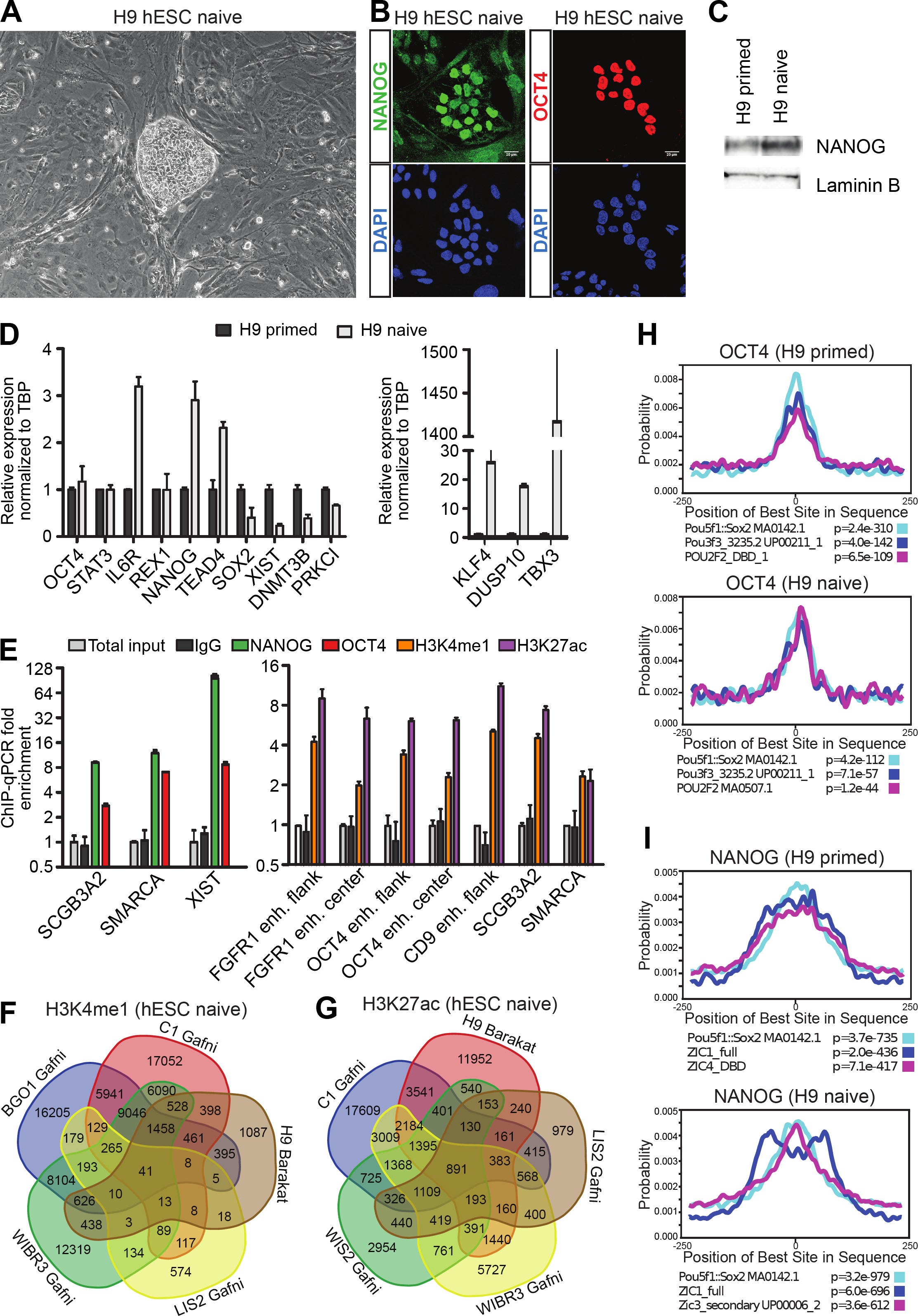
Conversion of primed to naive H9 human embryonic stem cells. **A**) Brightfield microscopy of a representative colony of H9 hESCs cultured in naive culture conditions on feeders for 10 passages showing a more dome-shaped colony morphology (compare to (Figure S1A). **B**) Immunofluorescence of naive H9 hESCs for NANOG (green) and OCT4 (red); DNA is stained with DAPI (blue). **C**) Immunoblot analysis of NANOG and LAMININ Bin primed and naive hESCs. **D**) qRT-PCR of pluripotency-related genes in H9 hESCs cultured in primed or naive conditions (10 passages). Error bars indicate standard deviations; n=3. **E**) ChIP-qPCR in naive H9 hESC with anti-NANOG, anti-OCT4 or rabbit IgG at three OCT4 and NANOG binding sites (from primed ChIP-seq data) in *SCGB3A2, SMARCA* (Kunarso et al., 2010) and *XIST* (left) or with anti-H3K4me1, anti-H3K27ac or rabbit IgG at binding sites near *FGFR1* (at central and flanking locations), *POU5F1* (at central and flanking locations), *CD9*, *SCGB3A2* and *SMARCA* (Rada-Iglesias et al., 2011) (right). The mean fold-enrichment is shown relative to total input control DNA, normalized to a non-bound site in *ACTB* (for NANOG and OCT4) or *NCAPD2* (for H3K4me1 and H3K27ac). Error bars indicate standard deviations; n= 3. **F-G**) Venn diagrams of the overlap between ChIP-seq peaks in naive hESCs, indicating the cell line and study for **F**)H3K4me1 and **G**) H3K27ac. The numbers indicate overlapping peaks. **H**) Local motif enrichment analysis using CentriMo (Bailey and Machanick, 2012) for OCT4 ChIP-seq data generated in this study for primed (upper panel) and naive H9 hESCs (lower panel). Top-3 identified motifs and their p-values are indicated. **I**) as **H,** but now for NANOG.

**Figure S6, related to Figure 4:**
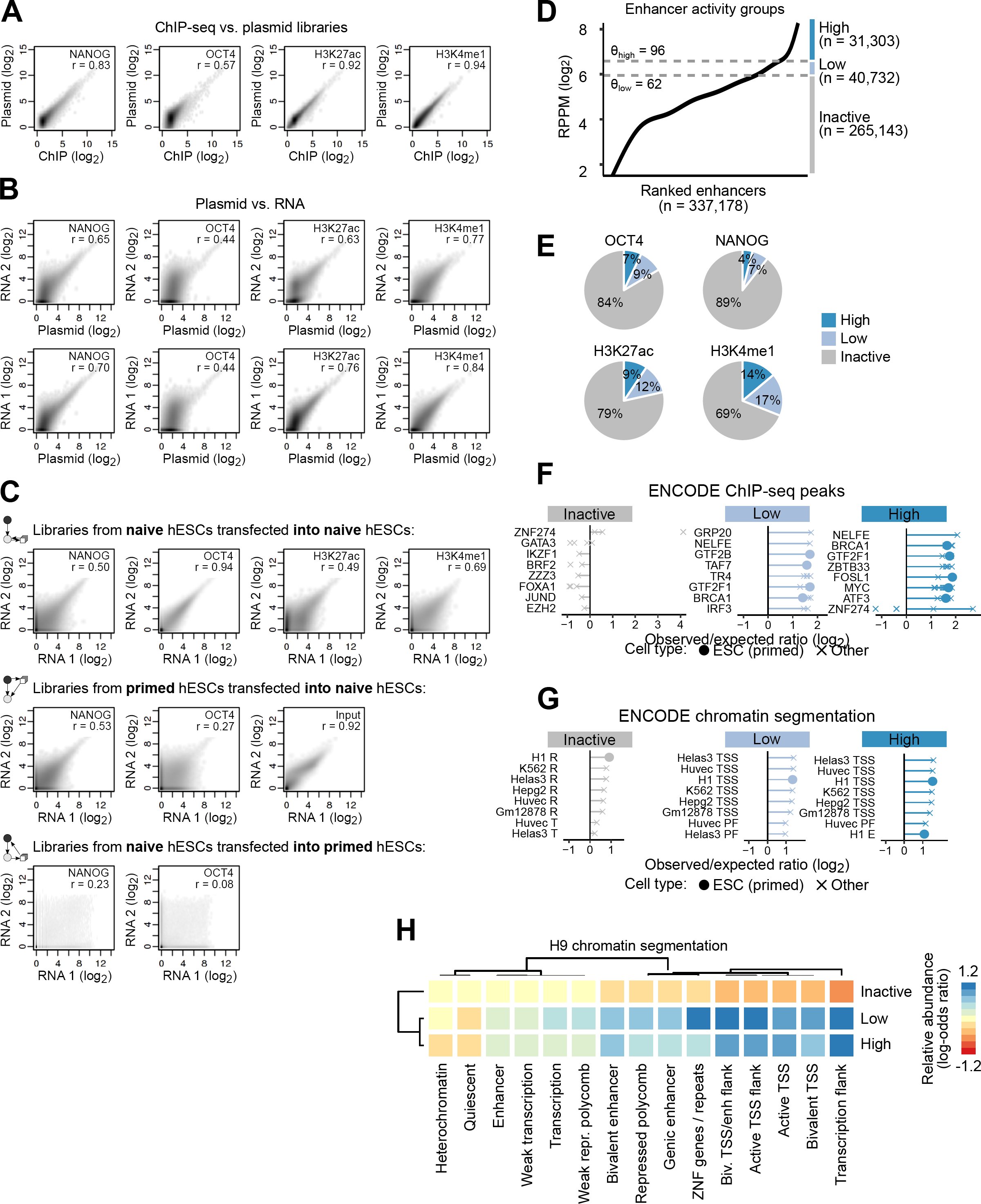
ChIP-STARR-seq in naive hESCs. **A-C**)Scatterplots contrasting normalized read counts (reads per million) in naive H9 hESCs per peak for different datasets. The Pearson correlation coefficient (*r*) for each comparison is indicated for each plot. **A**) Comparison of ChIP-seq datasets and the corresponding ChIP-STARR-seq plasmid library. **B**) Comparison between ChIP-STARR-seq plasmid libraries prior to transfection and the corresponding RNA-seq read counts generated from two replicates of GFP-positive naive H9 hESCs after transfection. **C**)Comparison between two replicates of GFP-positive naive H9 hESCs after transfection with libraries generated in naive H9 hESCs (top), with libraries generated in primed H9 hESCs (middle), or of GFP-positive primed H9 hESCs after transfection with libraries generated in naive H9 hESCs (bottom). (Zhang et al., 2008) **D**)Plot showing enhancer activity (ratio of ChIP-STARR RNA over plasmids; log_2_) in naive H9 hESCs ranked from lowest to highest across all measured enhancers (union of all peak calls). Three groups of enhancers were distinguished based on their activity and the thresholds (θ) are indicated in the plot by dashed lines. RPPM, reads per plasmid million. **E**) Relative distribution of highly active enhancers (RPPM ≥96), lowly active enhancers (RPPM 62-96) and inactive genomic regions (RPPM <62) in peaks called for the indicated factors. **F**) Relative enrichment of DNA-binding proteins (DBPs) from the ENCODE database (2012) in inactive genome regions, as well as lowly active and highly active enhancers in naive H9 hESCs (compare to Fig. 4B). Shown are the log_2_-odds ratios between observed percentages of enhancers overlapping binding sites of each given DBP in the respective groups over the percentage of overlaps in the entire enhancer dataset. Each dot represents one ChIP-seq dataset for the given DBP and the lines connect the most extreme dot with zero for visualization. For each category, the eight most enriched DBPs are shown ranked by their mean log-odds ratio. ChIP-seq datasets produced from hESCs are indicated as dots and those from other cell sources as crosses. Enrichments were calculated using LOLA (Sheffield and Bock, 2016). **G**) LOLA enrichment plots as in panel F, but showing instead the relative over-presentation of ENCODE chromatin segments from different cell lines in enhancers with different activity levels. E, enhancer; PF, promoter-flanking region; R, repressed; T, transcribed; TSS, transcription start site; WE, weak enhancer. **H**) Heatmap showing the relative enrichment of H9 chromatin segments from the Roadmap project (Kundaje et al., 2015) in regions with different levels of ChIP-STARR-seq activity from naïve hESCs. Heatmap colors report log-odds ratios. Rows and columns have been arranged by hierarchical clustering with complete linkage to put similar segment types together. TSS, transcription start site; enh, enhancer; ZNF, zinc-finger protein.

**Figure S7, related to Figure 4:**
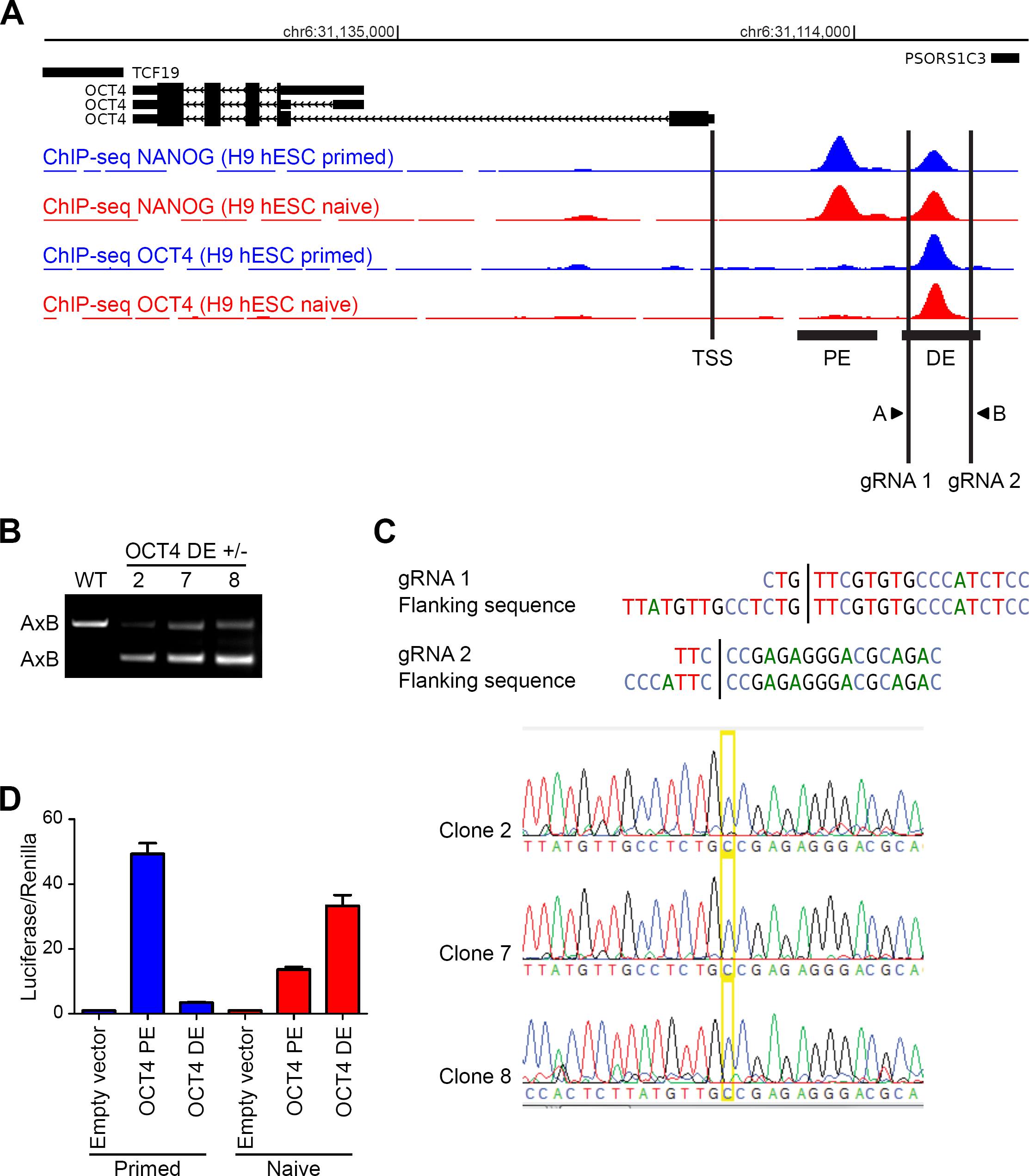
Generation of H9 hESCs cell lines with OCT4 distal enhancer deletion. **A**) Genome browser view of the *OCT4* (*Pou5F1*) locus and upstream region, with tracks from ChIP-seq for NANOG (blue) and OCT4 (red) in primed and naive H9 hESCs. The OCT4 transcriptional start site (TSS), proximal enhancer (PE), distal enhancer (DE) are indicated, as are the locations of gRNA1 and 2 used for DE deletion and genotyping primers A and B. Black bars represent sequences tested in luciferase assays. **B**) PCR genotyping using primers A and B confirms a 650 bp heterozygous deletion of the OCT4 DE in clones 2, 7 and 8. **C**) The sequences of gRNAs and flanking sequences are shown at the top. The line is the Cas9 cut site. Below is the dideoxy sequence traces of the deleted alleles in OCT4 DE+/- clones 2, 7 and 8, confirming expected sequence upon deletion. In the wt allele, the OCT4 DE sequence was still present (data not shown). **D**) Luciferase assay for proximal and distal OCT4 enhancer in primed (blue) and naive (red) H9 hESCs. Luciferase activity is reported as the fold enrichment in luciferase counts over empty vector, normalised to the Renilla transfection control. Error bars indicate standard deviations, n=2.

**Figure S8, related to Figure 6:**
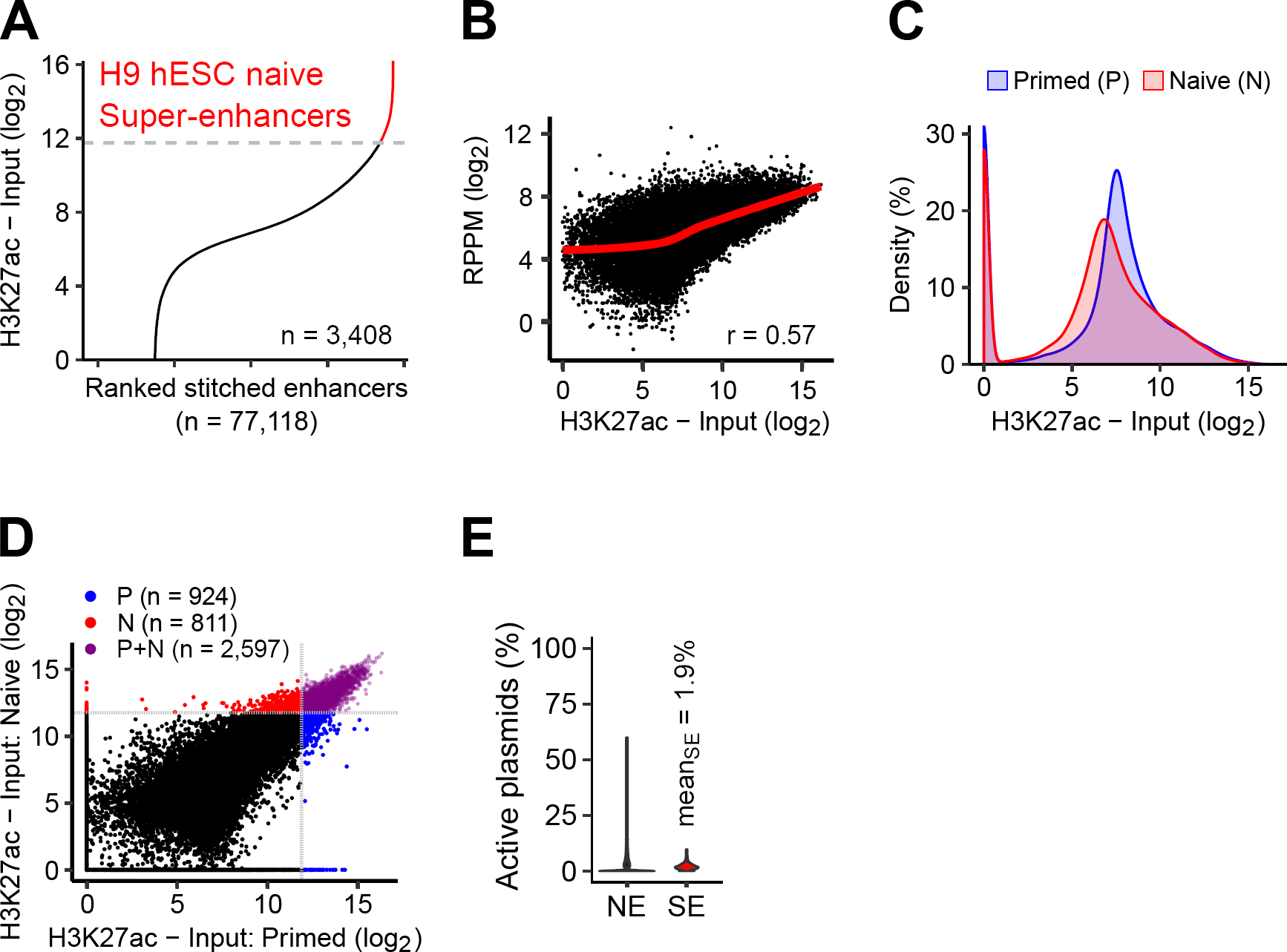
Super-enhancers and ChIP-STARR-seq in naive hESCs. **A**) Super-enhancers in naive H9 hESCs (SEs) were called from H3K27ac ChIP-seq data on enhancers stitched within 12.5kb windows using the ROSE software (Whyte et al., 2013). **B**)Scatterplot contrasting SE intensity in naive H9 hESCs (H3K27ac signal divided by input) with ChIP-STARR-seq activity. The Pearson correlation coefficient (*r*) is indicated and the red line represents a generalised additive model fit to the data. **C**) Kernel density plots showing the distribution of SE intensity values (H3K27ac signal divided by input) in primed and naive H9 hESCs. **D**) Scatterplot contrasting SE intensity values (H3K27ac signal divided by input) in primed and naive H9 hESCs. Regions called as SEs in primed, naive, or both hESCs are indicated in blue, red, or purple, respectively. **E**) Violin plots showing the proportion of active plasmids (RPPM ≥ 96) for 3,408 super-enhancers (SE) compared to normal enhancers (NE).

## Overview of Supplemental Tables

**Table S1 related to Figure 1: Data overview**

**Table S2 related to Figures 2, 3 and 4: LOLA and GREAT enrichments**

**Table S3 related to Figure 3: ESC and HK enhancer modules**

**Table S4 related to Figure 5: transposable elements observed/expected ratios**

**Table S5, related to Figures 1, 2, 4 and 6: Oligonucleotides used in this study**

**File S1 related to Figure 1, 2, 4 and 6: Genomic coordinates (BED files) of ChIP-seq peaks, ChIP-STARR-seq enhancer with activity level and of super-enhancers called in this study**

